# Spike N354 glycosylation augments SARS-CoV-2 fitness for human adaptation through multiple mechanisms

**DOI:** 10.1101/2024.01.29.577677

**Authors:** Pan Liu, Can Yue, Bo Meng, Tianhe Xiao, Sijie Yang, Shuo Liu, Fanchong Jian, Qianhui Zhu, Yuanling Yu, Yanyan Ren, Peng Wang, Yixin Li, Jinyue Wang, Xin Mao, Fei Shao, Youchun Wang, Ravindra Kumar Gupta, Yunlong Cao, Xiangxi Wang

## Abstract

Selective pressures have given rise to a number of SARS-CoV-2 variants during the prolonged course of the COVID-19 pandemic. Recently evolved variants differ from ancestors in additional glycosylation within the spike protein receptor-binding domain (RBD). Details of how the acquisition of glycosylation impacts viral fitness and human adaptation are not clearly understood. Here, we dissected the role of N354-linked glycosylation, acquired by BA.2.86 sub-lineages, as a RBD conformational control element in attenuating viral infectivity. The reduced infectivity could be recovered in the presence of heparin sulfate, which targets the “N354 pocket” to ease restrictions of conformational transition resulting in a “RBD-up” state, thereby conferring an adjustable infectivity. Furthermore, N354 glycosylation improved spike cleavage and cell-cell fusion, and in particular escaped one subset of ADCC antibodies. Together with reduced immunogenicity in hybrid immunity background, these indicate a single spike amino acid glycosylation event provides selective advantage in humans through multiple mechanisms.

**HIGHLIGHTS:** N354 glycosylation acts as a conformational control element to modulate infectivity Reduced infectivity could be recovered by altered binding mode of heparin sulfate N354 glycosylation improved fusogenicity and conferred escape from ADCC antibodies N354 glycosylation reduced immunogenicity and conferred immune evasion

## INTRODUCTION

The ongoing coronavirus disease 2019 (COVID-19) pandemic caused by severe acute respiratory syndrome coronavirus-2 (SARS-CoV-2) has lasted for nearly four years. A number of variants with improved fitness and immune evasion capabilities have been documented during the course of the pandemic ^1,2^. The emergence and circulation of Omicron represents a significant shift in the evolution trajectory of SARS-CoV-2 because this variant has over 30 mutations in its spike (S). Subsequently, several Omicron descendants, such as BA.2, BA.5, BQ.1 and XBB, have caused multiple waves of infections globally ^3,4^. The successive selection of these sublineages is primarily driven by immune pressure exerted by neutralizing antibodies present in human sera as a result of mass vaccinations or natural infections or breakthrough infections ^5^. However, immune evasion often comes at the cost of impairment in functionality and selection for antibody-escaping variants as well as accumulation of near-neutral mutations have led to suboptimal codon usage, thereby impacting functionality ^6^. Upon boosting with updated (Omicron-based) vaccine or single Omicron infection, immune responses to Omicron variants have been shown to be attenuated owning to the “original antigenic sin”. However, repeated Omicron exposures override ancestral SARS-CoV-2 immune imprinting, yielding high neutralizing titers against Omicron variants, including XBB sublineages ^7^. Given the extent of herd immunity raised by repeated Omicron exposures today, evolution of the virus by more nuanced human adaptation to overcome immune imprinting might be already under way.

Though decorated with fewer glycans than the HIV-1 Envelope protein, the dense glycan shield consisting of 22-23 *N*-glycosylation sequons per protomer is an essential feature of SARS-CoV-2 S architecture. The glycans have been shown to play intrinsic and extrinsic roles in protein folding, modulating conformational activation and immune evasion ^8,9^. Pathogenesis and selective sweeps analysis reveal that the evolution of glycosylation sites in SARS-CoV-2 S is intertwined with adaptive mutations of the amino acid sequence for successful cross-species transmission ^10^. For instance, loss of *N*-glycosylation at position 370 has been demonstrated to increase the receptor binding domain (RBD) in the up conformation, and thereby its exposure and accessibility for receptor recognition, improving viral infectivity in humans ^11,12^. Distinct from roles played by N370 glycosylation, many other glycans simply form a sugary barrier that shields antigenic epitopes vulnerable to neutralizing antibodies and immunogenic epitopes capable of eliciting neutralizing antibodies. Glycan shield density analysis reveals a strong correlation that viruses historically classified as “evasion strong” ^13^ had significantly elevated glycan shield densities ^14^. Consequently, sites of glycosylation are often positively selected during viral evolution in human host to increase glycan shield density. These assist the virus in evading the immune system, with impacts on infectivity ^15^. Therefore, acquisition of extra glycans that presumably improves viral fitness and adaptation in humans might have occurred over the long course of the SARS-CoV-2 pandemic.

Phylogenetic analysis of sarbecoviruses based on their S sequences reveals four clades: clade 1a (e.g. SARS-CoV-1), clade 1b (e.g. SARS-CoV-2), clade 2 (e.g. Rf1) and clade 3 (e.g. BtKY72) ^16^, among which coronaviruses from different clades display distinct clade-specific sequence characteristics at key sites shown to play roles in modulating viral infectivity, antigenicity and cross-species transmission (Figure 1A). In contrast to SARS-CoV-1, which emerged in 2002, was under control in 2003 and disappeared in 2004, SARS-CoV-2 seems to coexist with humans. After a prolonged period of nearly-complete global dominance of XBB subvariants, substantially mutated lineages, designated BA.2.86 sublineages, have quickly spread worldwide, out-competing XBB (Figure 1B) ^17–19^. BA.2.86 sublineages contain more than 30 mutations in the S when compared to XBB or its parental BA.2, and some of these mutations have been rarely observed in previously circulating variants (Figure 1C) ^17–19^. Surprisingly, Δ483, a key sequence feature of clade 1a viruses, has only recently been observed in SARS-CoV-2 variants. The substitution P621S is also a feature of SARS-CoV-1 variants. P681R is a fusion-enhancing modulator contained in Delta. Both P621S and P681R have been selected in the BA.2.86 lineage (Figures 1A and S1A). The mutation K356T, predicted to acquire glycosylation at N354 due to the formation of a standard N-linked glycosylation site motif (NXT/S) occurred only in recently emerging SARS-CoV-2 variants, rather than in early SARS-CoV-2 variants and sarbecoviruses from other clades (Figures 1A and S1A). In addition, the new substitution of H245N in BA.2.86 yields one extra glycan at N245, further suggesting a gradual accumulation of a glycan shield. Coincidentally, a distinct footprint of positive selection located around a new non-synonymous change (A1067C; K356T) within the RBD was found through scanning over 180,000 SARS-CoV-2 genomes deposited from 1^st^ Sep 2023 to 1^st^ Jan 2024, indicating a selective sweep (Figure 1D). Details of how the acquisition of glycosylation sites impacts the fitness of the virus are not clearly understood.

**Figure 1:**
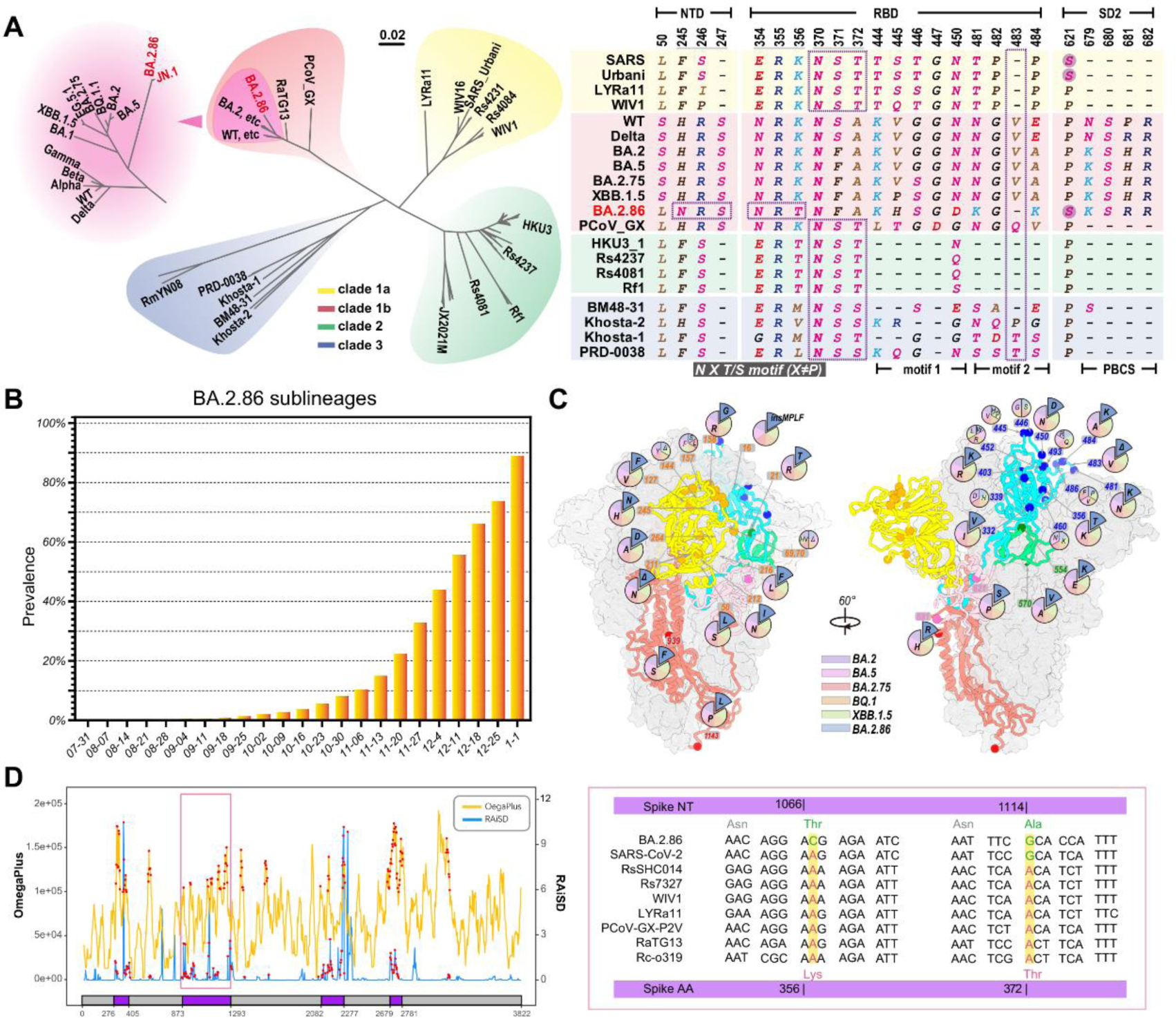
Selective advantages in the Spike gene. (A) Phylogenetic tree of sarbecoviruses based on S sequences and sequence features of S protein of selected variants from four clades. At the left panel, four clades are shadowed in yellow (clade 1a), red (clade 1b), green (clade 2) and blue (clade 3), respectively. For clade 1b, lineages of SARS-CoV-2 are zoomed in. Some typical variants are labeled and BA.2.86 and JN.1 are highlighted. At the right panel, motifs satisfying N-glycosylation (NXS/T, X≠P) are circled and residue 483 and 621 on S are highlighted. (B) Bar chart of the relative prevalence of BA.2.86 sublineages as of the first week of 2024. (C) Locations and residue diversities of mutations carried by BA.2.86 spike. Only residues on S differing from BA.2 are selected. Mutations unique to BA.2.86 are highlighted by larger detached sector diagram. For color scheme, NTD, RBD, SD1, SD2 and S2 are colored in yellow, cyan, green, pink and light red. The CA atoms of mutated residues are shown as spheres. (D) Selective sweep regions (shown as purple blocks) identified in SARS-CoV-2 genomes using OmegaPlus (yellow lines) and RAiSD (blue lines). Important non-synonymous differences are highlighted at the right panel.

## Results

### N354 glycosylation modulates RBD conformation

To explore the putative acquisition of glycosylation at N245 and N354 in more recent variants, we determined the asymmetric cryo-EM reconstructions of the BA.2.86 and JN.1 S-trimer at pH 7.4, to mimic the physiological conditions, at atomic resolution (Figures 2A, S2A, S2B and Table S1). In contrast to S-trimers from most variants ranging from WT, D614G through Alpha, Delta, Omicron to XBB and XBB.1.5, which sample the RBD-up conformation more frequently (>50%), BA.2.86 and JN.1 S-trimers dominantly adopt a closed state with all three RBDs in the down configuration (Figure 2A), similar to structural observations of VAS5, a highly attenuated SARS-CoV-2 vaccine candidate ^20^. In line with these structural observations, BA.2.86 was previously reported to have compromised infectivity and attenuated pathogenicity in animal models ^21,22^. Compared to other variants, there are two additional glycosylation related modifications at N245 and N354 in BA.2.86 sublineages, among which N245 glycan lies at outermost region of each NTD around the triangular vertices of the S-trimer (Figure 2B). Notably, the N354 glycan resides in a cleft formed by the NTD and RBD from two neighboring subunits, and establishes hydrogen bonds with T167 of NTD and with E340 of RBD, respectively, acting like a “bolt” to lock the S-trimer in “RBD-down” state (Figures 2B and 2C). This is akin to roles played by LA, a polyunsaturated fatty acid found in the RBD in stabilizing the RBD-down state by locking the conformation of the S-trimer ^23^. In line with this, the N354 glycosylation confers a more compact architecture in the region formed by the three NTDs and RBDs (Figure 2C). When the closed S-trimers were superimposed with its counterparts from XBB.1.5, the NTD, RBD and SD1 from BA.2.86 moved inward to the three-fold axis with the shift distances of 6Å, 3Å and 2Å, respectively (Figure 2D), forming a tight packing between NTD and RBD.

**Figure 2:**
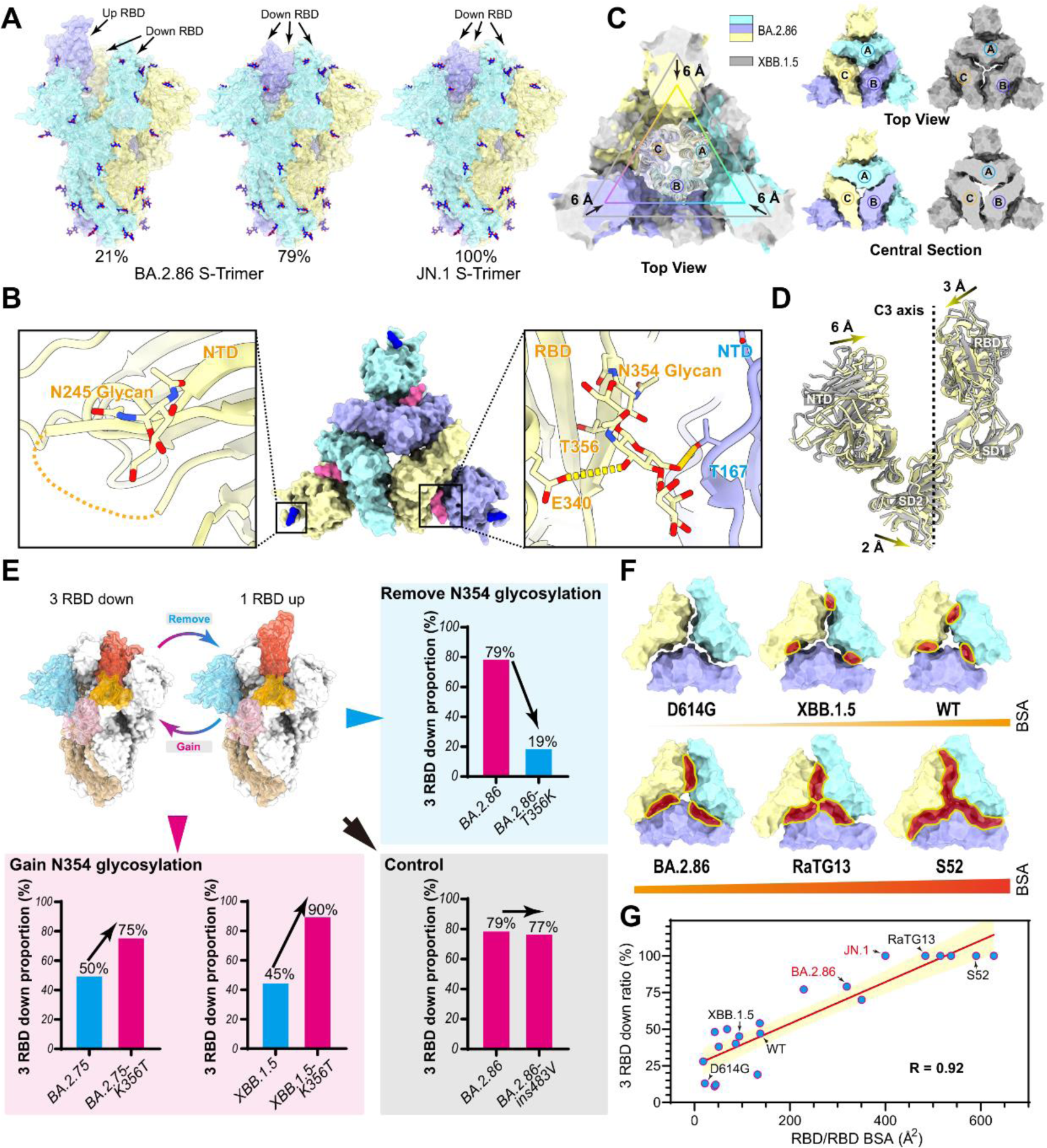
N354 glycosylation modulates RBD conformation. (A) Surface characterization of S-trimer of BA.2.86 and JN.1 in “up” and “closed” conformational states. The three subunits of S protein were colored in yellow, light blue, and purple respectively, and the N-glycans were highlighted with sticks. (B) The glycosylation modifications at N245 and N354 in BA.2.86 sublines were shown in detail. (C) RBD of the “closed” conformation of BA.2.86 S-trimer was superimposed with the XBB.1.5 S-trimer (color scheme same as Figure 2A). Top view (top right) and center section (bottom right) show intersubunit contacts of BA.2.86 and XBB.1.5 S-trimers. (D) The conformational change details between BA.2.86 (yellow) and XBB.1.5 (gray) in S1 subunit. The shift distances and directions of NTD, RBD and SD2 towards the 3-fold axis are labeled. (E) The role of N354 glycosylation in regulating changes in the “up” and “down” conformational ratio of the RBD. Three groups (“Remove N354 glycosylation”, “Gain N354 glycosylation” and “Control”) are located in the upper right corner, the lower left corner and lower right conner, respectively. In each group, the proportion of “RBD down” conformation are displayed with bar chart. Blue and pink bars represent variants without and with N354 glycosylation. (F) The contact areas between RBDs of D614G, XBB.1.5, WT, BA.2.86, RaTG13, and S52 were compared. (G) A correlation plot was created between the contact area between RBD subunits and the “RBD-down” rate.

To further verify the role of N354 glycosylation in modulating RBD up/down disposition, four additional modified constructs, BA.2.86-T356K, predicted to remove glycosylation at N354, XBB.1.5-K356T and BA.2.75-K356T, predicted to acquire N354 glycan, together with BA.2.86-ins483V as a control, were characterized, and their structural features were compared with the ancestral strain (Figures 2E and S2C). We observed N354 glycosylation dramatically increased the proportion of the “closed” form from 19% to 79% in BA.2.86, 50% to 75% in BA.2.75 and 45% to 90% in XBB.1.5, making the ‘closed’ form being the dominant population (Figure 2E). However, the insertion of V483 had little effect on the modulation of the RBD conformation (Figure 2E), suggesting the ‘closed’ and ‘open’ form transition was not due to the general effects of mutations in RBD but the specific presence of N354 glycosylation. Results of our studies together with previous studies on N-glycans at N165, N234 and N370 (not found in all SARS-CoV-2 variants) capable of participating in RBD up/down disposition ^8,11,24^, allows us to propose detailed molecular basis for RBD conformation modulation, in which compact inter-subunit (S1/S1) arrangements relay a cascade of interactions mediated by specific N-glycans via tight connections with neighboring subunits or intrinsic packing modes, facilitating the RBD-down switch (Figure 2F). To further decipher the relationship between inter-subunit contacts and “RBD-down” rate, we systematically analyzed the S1/S1 or RBD/RBD or NTD-RBD/NTD-RBD interactions and calculated the “RBD-down” rates of available SARS-CoV-2 S-trimer structures (n=21), including ours in this study. We found that contact areas between RBDs determine the “RBD-down” rate with a compelling correlation of 0.92 (Figures 2F and S2D). Of note, RBD/RBD contact areas of over 400 Å^2^ drive the S-trimer to be in the closed state only, which is a common feature in animal derived sarbecoviruses, such as bat RaTG13, pangolin PCoV_GX and BANAL-20-52. However, those sarbecoviruses are able to bind ACE2, but with decreased infectivity in human cells ^25–27^. These results promoted us to hypotheses that the N354 glycan, nestled between the NTD and RBD interface may function as a conformational control element for modulating infectivity.

## N354 glycosylation decreases infectivity irrespective of comparable hACE2 binding

Given the fact that the presence of the N354-linked glycan favors the closed state in BA.2.86 sublineages, this presumably leads to a compromised infectivity and attenuated pathogenicity. To verify this, we first compared the infectivity of BA.2.86 and representative SARS-CoV-2 variants by using pseudotyped viruses in HEK293T cells overexpressing hACE2 (293T-ACE2) or TMPRESS2 (293T-TMPRESS2) or both hACE2 and TMPRESS2 (293T-ACE2-TMPRESS2) and in widely used cell lines, such as Vero, H1299, Huh-7 and Calu-3. Like Omicron variants, BA.2.86 can enter cells via endosomes as well as through TMPRSS2 but prefers ACE2-mediated infection (Figure S3A) in concordance with authentic BA.2.86 infection results ^28^. Overall BA.2.86 exhibited a partially decreased infectivity compared to most Omicron variants (Figures 3A and S3A), which largely matches with recent studies ^17–19^, albeit improved entry in lung cells rather than other cells relative to specific variants being observed as well ^29,30^. These *in vitro* findings correlate to *in vivo* clinical observations that currently there are no reports of elevated disease severity associated with this variant ^31^. Virus-host receptor engagement and membrane fusion directly affect viral infection efficiency. To further investigate if the RBD-dynamics modulator (N354 glycosylation) and putative fusion-related mutation (P621S) impact infectivity, we constructed BA.2/XBB.1.5/BA.2.86 derivatives that bear respective mutations and measured their infectivity in 293T-ACE2, Vero and Huh-7 cells (Figure 3B). As expected, acquisition of N354 glycosylation generated by the K356T mutation in BA.2 and XBB.1.5 decreased their infectivity and loss of N354 glycosylation raised by the reverse mutation T356K in BA.2.86 increased its infectivity (Figure 3B). Surprisingly, the substitution of P621S predicted to affect fusion activity in BA.2 and XBB.1.5 contributed to the increased infectivity; in turn the reverse substitution of S621P in BA.2.86 resulted in a further decreased infectivity (Figure 3B). Coincidentally, the N354 glycosylation (K356T mutation) first emerged in BA.2.75.5 and then in XBB.1.5.44, but they did not display growth advantages compared to the prevalent variants, presumably due to the dramatically reduced infectivity (Figure S1A). S621P to a large extent compensated the decreased infectivity conferred by the N354 glycosylation.

**Figure 3:**
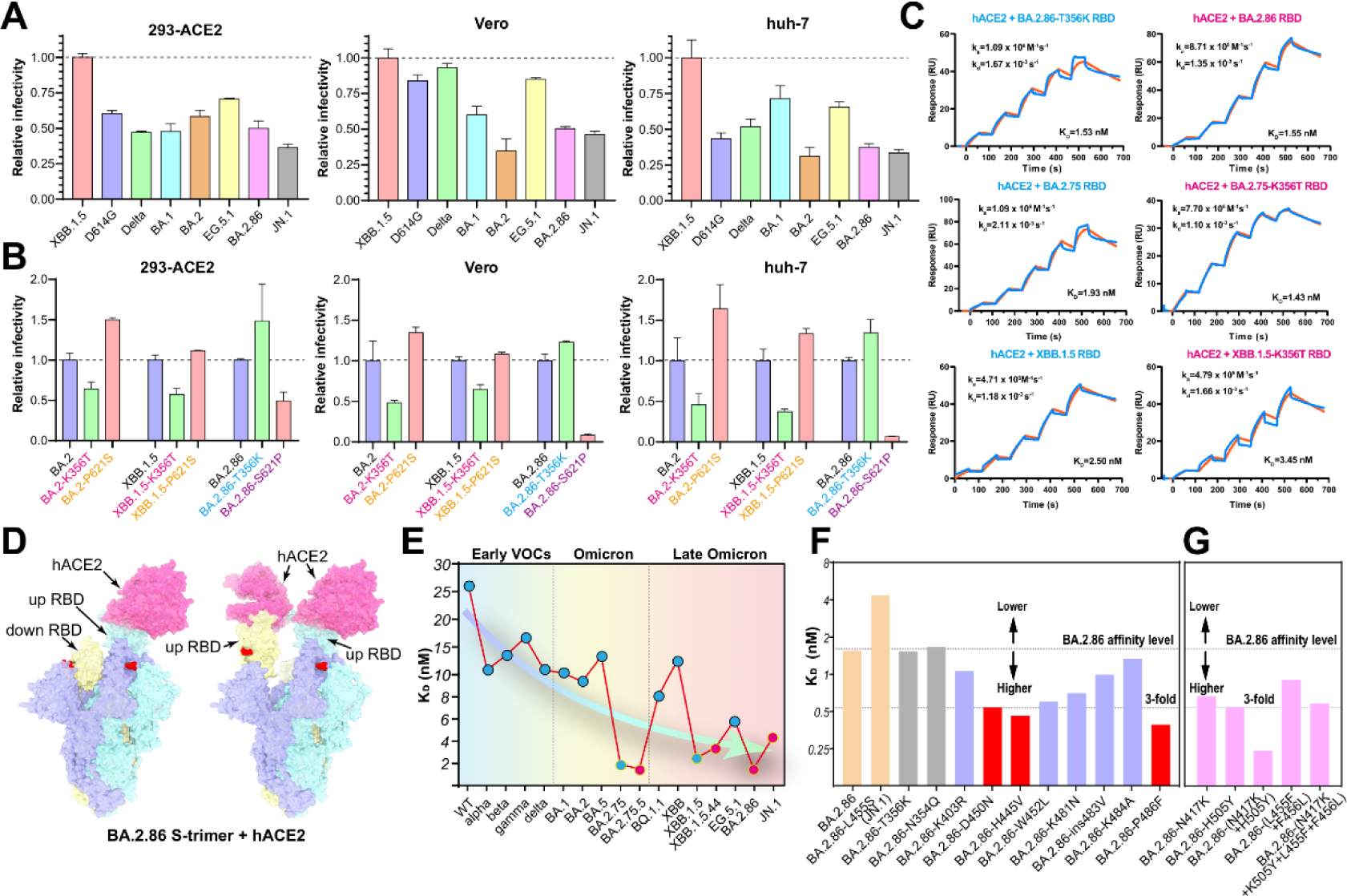
N354 glycosylation decreases infectivity and binding affinity to hACE2. (A) Relative infectivity of XBB.1.5, D614G, Delta BA.1, BA.2, EG.5.1, BA.2.86 and JN.1. Vesicular stomatitis virus-based pseudoviruses were used to test the efficiency of infecting 293T-ACE2, Vero, Huh-7 cells. Error bars represent the mean ± SD of three replicates. All raw data of infectivity are normalized by XBB.1.5. (B) Relative infectivity of BA.2, XBB.1.5, and BA.2.86 variants with mutations at positions 356 and 621, compared to their respective wild types, was evaluated in 293T-ACE2, Vero, and Huh-7 cells. This evaluation was done using a pseudovirus system based on the vesicular stomatitis virus. Error bars represent the mean ± SD of three replicates. (C) The impact of glycosylation at position 356 of BA.2.86, BA.2.75, and XBB.1.5 RBDs on the binding affinity to hACE2 was assessed by SPR. (D) Surface characterization two “up” RBD conformations of BA.2.86 S-trimer binding to hACE2 determined by Cryo-EM. The color scheme for three subunits of S are consistent with figure 2A and hACE2 is colored in pink. (E) Changes in affinity of binding hACE2 from early SARS-CoV-2 variants of concern (VOCs) and omicron variants to late omicron variants. (F) The effect of a single substitution on the binding affinity to hACE2 was assessed using SPR. Mutations that greatly enhance, moderately enhance and decrease the affinity to hACE2 are indicated in red, light purple and yellow. The cutoff value of greatly increasing affinity is set as 3-fold change in K_D_ value relative to BA.2.86. (G) Evaluation of binding affinity to hACE2 of the variants with 1-2 mutations on the RBM of BA.2.86 by SPR. These variants are based on predictions of increased binding affinity to hACE2.

We next sought to examine the possibility that the N354 glycosylation mediated impaired infectivity may be related to decreased binding affinity to hACE2. For this, three paired groups of six variants (BA.2.75/BA.2.75.5; XBB.1.5/XBB.1.5.44; BA.2.86/BA.2.86-T356K) were carefully selected due to the only difference being the N354 glycosylation or not in their RBDs. Surface plasmon resonance (SPR) results demonstrated that RBDs with or without the N354 glycosylation showed comparable binding affinities to hACE2, indicating that the N354 glycosylation does not impact hACE2 binding (Figure 3C), despite N354 glycosylation promoting the S-trimer “RBD down” state. To further structurally verify this, we determined the cryo-EM structure of the BA.2.86 S-trimer in complex with hACE2 (Figures 3D and S3B; Table S1). Like most complex structures, one or two copies of hACE2 are bound to the RBDs in the up configuration (Figure 3D). Consistent with binding results, both the N354 glycan and T356 are located far away from the interface (Figure S3C). However, we noted that variants with the N354 glycosylation exhibited very high affinities to hACE2, reflecting that tight binding might be required to further recover the compromised infectivity (Figure 3E). To further explore the contribution on hACE2 tight binding exemplified by BA.2.86, we evaluated the individual substitution (including reverse mutation) of N354Q, T356K, K403R, D450N, H445V, W452L, L455S, K481N, ins483V, K484A or P486F in BA.2.86 RBD on the hACE2 affinity. Surprisingly, all single mutations except for L455S identified in JN.1 displayed, to some extent, increased binding affinities and single reverse mutation of D450N, H445V and P486F induced an approximately 3-fold affinity increase (Figure 3F). Furthermore, mutations identified in key variants were also evaluated, among which a reverse-mutated combination of N417K and H505Y, as well as a pair of Flip-mutations of L455F and F456L synergistically enhanced hACE2 binding (Figure 3G). Structural comparisons revealed that the substitutions of N417K and Y505 established extra hydrogen bonds with D30 and E37 on ACE2, and mutations, including P486F, H445V, W452L and L455F-F456L augmented hydrophilic interactions of microenvironment, increasing the binding capabilities (Figure S3D). These suggest that successful selection for acquisition of the N354 glycosylation possibly needs to be accompanied by tighter ACE2 binding together with the P621S substitution, co-manipulating the infectivity.

### Decreased infectivity by N354 glycosylation can be restored by HS

Viruses like influenza and coronavirus use glycans as entry factors. In particular, the initial interaction with host cells is mediated by glycans ^32,33^. Growing evidence supports a role for negatively charged glycans, such as heparin sulfate (HS) as entry co-factors for SARS-CoV-2 ^34^. More importantly, these entry co-factors and furin expression are specially more abundant in nasal epithelial cells and upper airway cells compared to those in lungs ^35^. Perhaps correlated with this, Omicron variants display gradual increase in binding affinity to HS compared to early VOCs ^36^, presumably leading to a tropism alteration during SARS-CoV-2 evolution. Together with increased positive charges (Figure S4A) and nearly all closed S-trimers mediated by the N354 glycosylation (Figure 2A), these raise a possibility of altered entry factor usage in nasal epithelial and upper airway microenvironments. To mimic authentic virus infection at multiple steps of viral life cycles in nasal and upper airway tracks, we evaluated co-factor usage efficiency in representative variants via pre-treatment of virus-like particles (SC2-VLPs) ^37^ with various concentrations of free HS prior to infection by using 293T-ACE2-furin cells (Figure 4A). We observed that free HS displayed a dose-dependent reduced infection of BA.5, BA.2.75 and XBB.1.5, consistent with previously reported inhibition in S binding and infection by authentic SARS-CoV-2 ^34^. Surprisingly, HS treatment dramatically increased BA.2.86 infection, exceeding XBB.1.5 infectivity (Figure 4A), which suggests that abundant HS and furin aided the recovery of the decreased infectivity for BA.2.86. To further decipher the underlying mechanism, we constructed BA.5-K356T, BA.2.75-K356T, XBB.1.5-K356T and BA.2.86-T356K SC2-VLPs to gain or remove the N354 glycosylation, respectively, and compared the effect in HS treatment with their parental SC2-VLPs. Strikingly, all the N354 glycosylated SC2-VLPs exhibited a dose-dependent enhanced infectivity upon HS treatment, reaching up to the infectivity level for their parental variants and loss of the N354 glycosylation largely eliminated BA.2.86 HS-treated infectious enhancement (Figure 4A), indicative of differential usage of HS as a cofactor for modulating infectivity of the N354 glycosylated variants in furin/hACE2 enriched microenvironments, which has recently been reflected by experimental observations of potent infections for BA.2.86 in nasal epithelial cells ^38^. In line with these results, the N354 glycosylation partially impaired binding of HS to RBDs (Figure 4B). Intriguingly, HS mediated enhancements of infectivity for the N354 glycosylated variants became marginal by using pseudo-typed viruses in 293T-ACE2 cells, while HS dose-dependent inhibitions of infectivity for variants without the N354 glycosylation were still straightforward (Figure S4B). The possible reason for differences yielded from two systems might lie in excessive redundancy of spikes decorated on VSV-based pseudoviruses, in which limited numbers of the “open” spikes can initialize a successful infection even though majority in the closed state, largely diluting roles played by HS in modulating infectivity of the N354 glycosylated variants via promoting the RBD-up transition. Collectively, these revealed that HS and furin enriched microenvironment might offset the impaired infectivity caused by the N354 glycosylation and even possibly support the shift in tropism towards HS-abundant cells.

**Figure 4:**
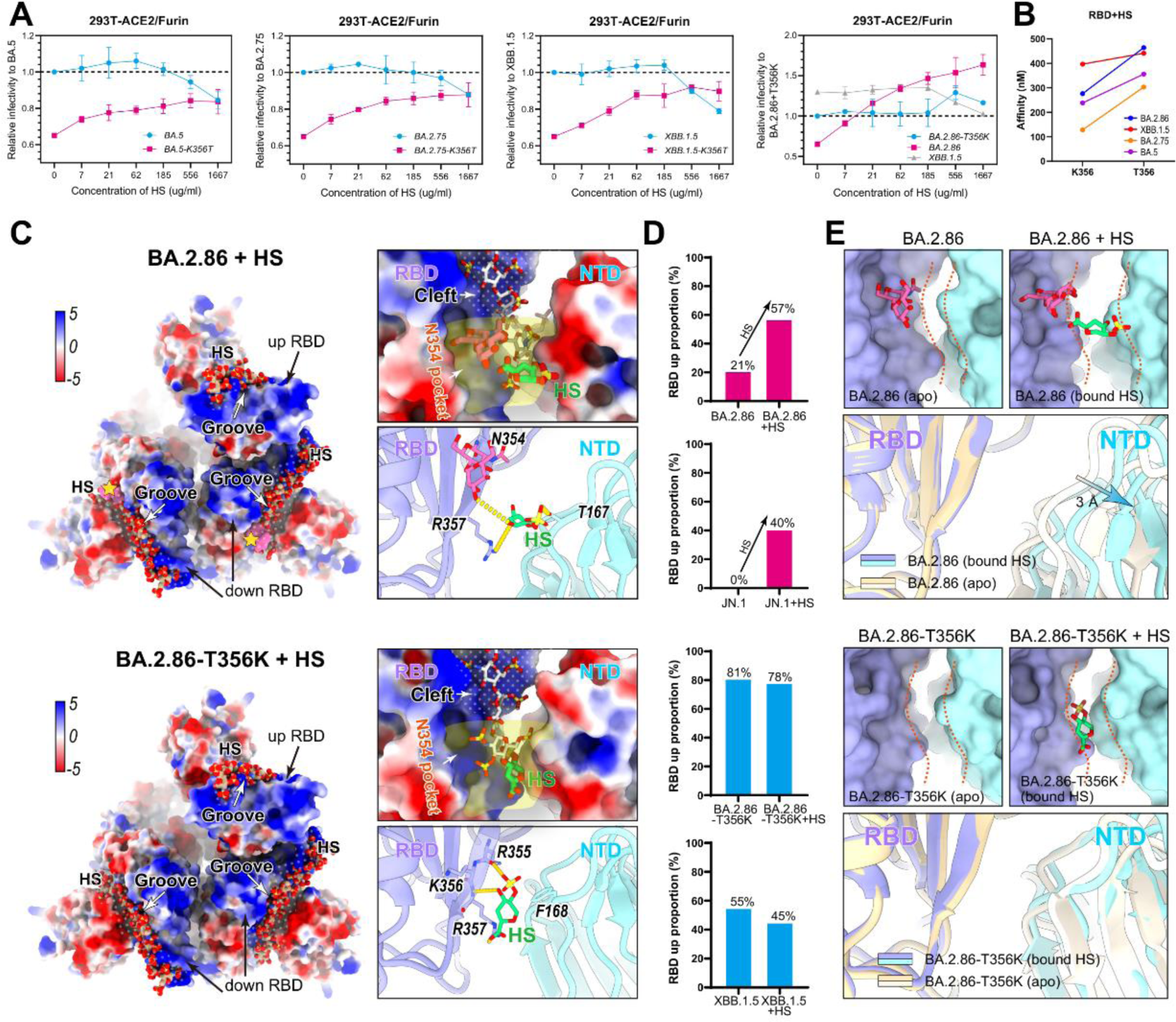
Mechanism of the ability of heparan sulfate recovering decreased infectivity by N354 glycosylation. (A) The ability of BA.5, BA.2.75, XBB.1.5, BA.2.86-T356K (blue) and their corresponding K356T (pink) mutant virus-like particles (SC2-VLPs) to infect HEK293T cells overexpressing ACE2 and Furin (293T-ACE2/Furin) treated with various concentrations of free heparin sulfate (HS). (B) Binding affinity of RBDs of BA.5, BA.2.75, XBB.1.5, BA.2.86-T356K and their corresponding K356T mutant to HS tested by SPR. (C) The Cryo-EM structures of BA.2.86 and BA.2.86-T356K S-trimer bound to HS are showed on the upper and lower panels, respectively. In each panel’s left corner, HS was docked to S-trimer by MOE. The binding grooves of HS are indicated by “dotted zones” on the electrostatic surface of S-trimer. pink surface of N354 glycosylation on BA.2.86 is highlighted by yellow stars. On the upper right corner, the explicit binding location of HS “N354 pocket” in groove is zoomed in and indicated by light yellow shadow. HS determined by Cryo-EM and docked by MOE are colored in green and white, respectively. On the lower right corner, interface details of HS determined by Cryo-EM with S-trimer are shown. Hydrogen bonds are displayed by yellow dashed lines. The unit of the value for color bar is kcal/(mol·e) at 298 K. (D) Influence of co-incubation with HS on the “RBD-up” conformational proportion within the S-trimer of BA.2.86, JN.1, BA.2.86-T356K, and XBB.1.5. BA.2.86 and JN.1, which are glycosylated at position N354, are represented by red bars, while BA.2.86-T356K and XBB.1.5, which lack glycosylation at position N354, are represented by blue bars. (E) Surface and cartoon representation of HS binding grooves consisting of a pair of spatially adjacent RBD (purple) and NTD (cyan) from different subunits for BA.2.86 (upper panel) and BA.2.86-T356K (lower panel). In each panel, apo state and bound HS state are showed on the left and right. Distance of HS binding grooves is indicated by orange dashed curves.

To understand SARS-CoV-2 engagement of HS cofactor and how the N354 glycan alters HS usage at molecular level, we determined cryo-EM structures of XBB.1.5, BA.2.86, JN.1 and BA.2.86-T356K in complex with HS at 3.2–3.8 Å (Figures 4C and S4C). Interestingly, incubation with HS led to marked conformational alteration, yielding substantially increased “RBD-up” state in the N354 glycosylated S-trimers, but had limited impact in RBD conformation modulation for S-trimers without the N354 glycosylation (Figure 4D). Due to structural heterogeneity and flexibility of HS, only the density for the part of HS basic unit, IdoA(2S) (2-O-sulfo-α-L-iduronic acid) is straightforward, allowing identification of the location of major binding site and detailed interactions (Figure 4C). In contrast to binding of sialoglycan to the domain A (corresponding to the NTD in SARS-CoVs) in HKU1 and MERS-CoV ^39,40^, HS mainly targets a semi-open, shallow, elongated cavity composed of a number of positively charged residues on RBD, downstream within a deep groove, named the N354 pocket, constructed by the residues N354, R355, K356T and R357 from RBD and T157 and F168 from neighboring NTD, is occupied by the HS fragment (Figure 4C). The HS fragment is poised to directly interact with R355 and R357 through hydrogen bonds and a salt bridge, meanwhile residues K356, N354, R346 and R466 might contribute to further coordinate the oligosaccharide (Figure 4C). Notably, the absence of the N354 glycan in the immediate vicinity of the binding groove probably facilitates unobstructed engagement of HS, in line with the observed affinity; however, the presence of the N354 glycan together with the bound HS widens the binding groove by 3 Å, pushing the neighboring NTD outwards and thereby conferring a relatively relaxed upper arrangement (Figure 4E). The high proportion of the “3-RBD-down” state led by the N354 glycan mediated compact upper architecture could be partially converted to the “RBD-up” state upon HS binding, which explained the experimental observation that HS treatment increased infectivity for the N354 glycosylated variants.

### N354 glycosylation affects S cleavage and fusogenicity

We next sought to examine the possibility that the impaired infectivity caused by the N354 glycosylation in some cells might be related to differential S cleavage. For instance, Delta, which is known to show higher infectivity, is associated with a highly cleaved S protein and more efficient TMPRSS2 usage for entry ^41^. Furin cleavage dependent on the polybasic cleavage site (PBCS) between S1 and S2 is a key step in regulating virus infectivity and fusion activity ^41–43^. Alterations at P681 in PBCS have been observed in multiple SARS-CoV-2 lineages, H681 in Alpha and most Omicron variants; R681 in Delta and BA.2.86 (Figure 5A). To evaluate the cleavage efficiency, we first tested the cells used for WT, BA.2 and BA.2.86 pseudoviruses production by western blot analysis. We found substantially improved cleavage in BA.2.86 compared with BA.2 as evidenced by the ratio of S1/S2 to full-length S, despite slightly lower than that in WT and Delta (Figures 5B and 5C), suggesting that mutation at P681 contributes non-exclusively to S cleavage. To further investigate putative contribution on enhanced cleavage, we also evaluated S-cleavage in BA.2.86-T356K and BA.2.86-S621P. Interestingly, the loss of N354 glycosylation dramatically through T356K mutation decreased cleavage efficiency and the reversion of S621P moderately increased S cleavage (Figure 5C), indicative of BA.2.86 S-trimers being more likely in a postfusion conformation under furin enriched microenvironments. Given our data showing inefficient TMPRSS2 usage for BA.2.86 sublineages, N354 glycosylation appears to contribute to the negative correlation between cleavage efficiency and infectivity (Figures 3B and 5C).

**Figure 5:**
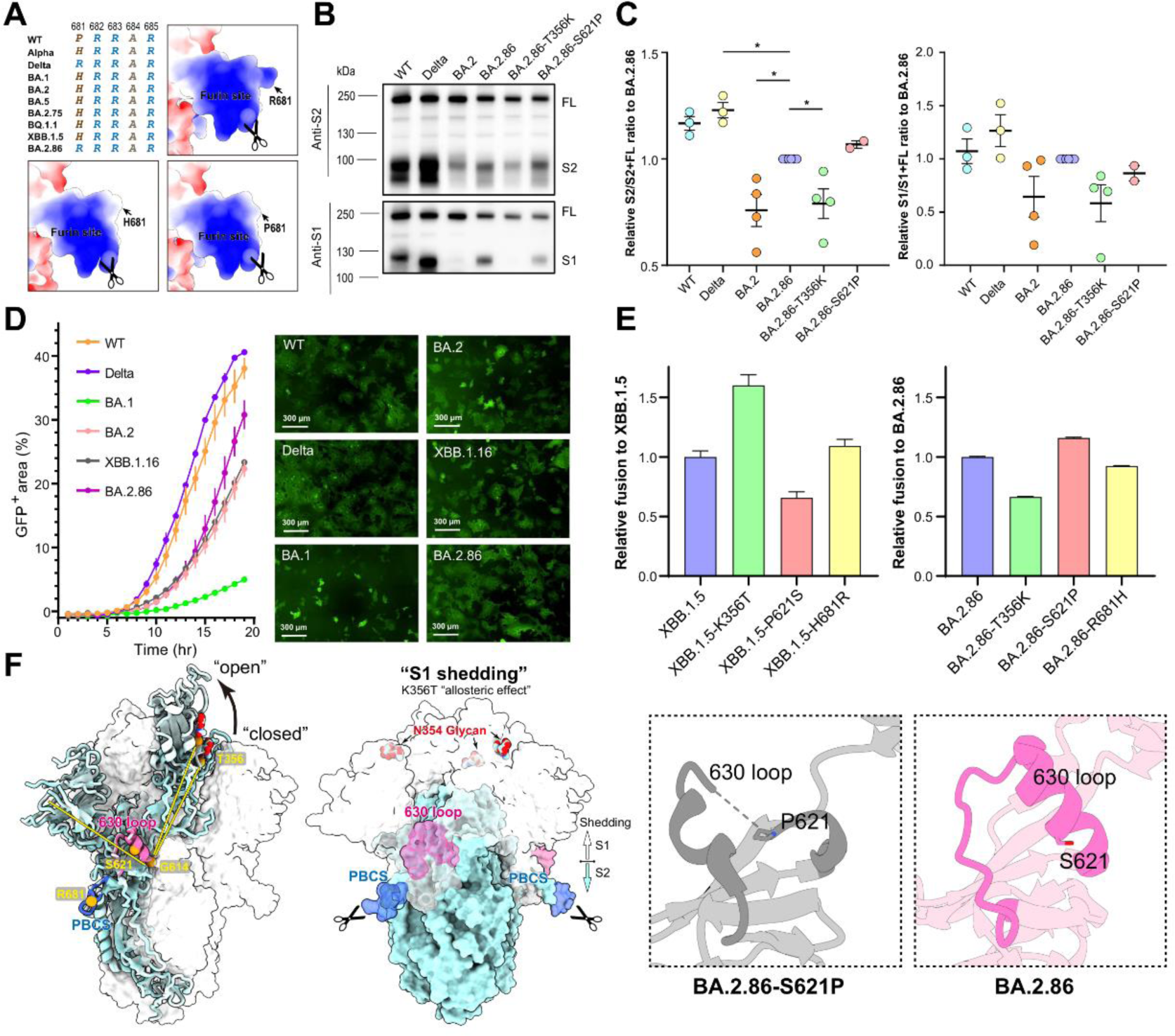
S cleavage and fusogenicity of BA.2.86. (A) Sequence alignment and modeled charge surface representation of PBCS of P681, H681 and R681 SARS-CoV-2 variants. S cleavage efficiency evaluated by Western blotting (B) and image grayscale analysis (C) for WT, Delta, BA.2, BA.2.86, BA.2.86-T356K and BA.2.86-S621P. (D) Fusogenicity comparation among WT, Delta, BA.1, BA.2, XBB.1.16 and BA.2.86 by using a split GFP system. (E) Fusogenicity of variants with K356T, P621S and H681R mutation on XBB.1.5 relative to XBB.1.5 (left) and variants with T356K, S621P and R681H mutation on BA.2.86 relative to BA.2.86 (right). (F) The location of N354 glycan (red), 630 loop (pink) and PBCS (blue) on S-trimer is shown on the left. T356, G614, S621 and R681 are displayed as orange spheres. Pattern diagram of K356T increasing S1 shedding by allosteric effect is shown in the middle. Cartoon representations of 630 loop of BA.2.86-S621P and BA.2.86 are zoomed in on the right.

The ability of SARS-CoV-2 S to induce cell–cell fusion that could provide an additional route for viral dissemination and promote immune evasion correlates with the PBCS, S cleavage efficiency and the usage of TMPRSS2 ^41^. Given the requirement of TMPRSS2 and S cleavage for optimal cell–cell fusion, Delta displayed the highest fusion activity; on the contrary BA.1 had quite low fusogenicity ^42,43^. We hypothesized based on the increased cleavage efficiency the fusion efficiency is altered in BA.2.86 in comparison to BA.2. To examine this, we used a split GFP system ^41^ to monitor cell–cell fusion in real time. We observed that BA.2.86 showed an increment in cell-cell fusion compared to BA.2, but was still demonstrably lower than WT and Delta (Figure 5D). The efficiency in fusion is reversely correlated with the stabilities of S-trimers (Figure S5A), which can be explained by the fact that structural transitions from the prefusion to the postfusion stage involve a series of conformational changes between domains and subunits, a prerequisite for viral fusion. Structural comparisons with BA.1 revealed reduced interactions between domains, including NTD-RBD, RBD-SD1/SD1-S2 and S2-S2 in BA.2.86, structurally explaining compromised stability (Figure S5B). Not surprisingly, either the loss of the N354 glycan or substitution R681P/H in BA.2.86 substantially reduced the cell-cell fusion activity; on the contrary acquisition of the N354 glycan or the mutation H681R based on XBB.1.5 contributed to increased fusion activity (Figure 5E). The improved S processing and fusion might be related to the structural observation that the N354 glycan tightly cements the NTD and RBD from adjacent subunits together presumably aiding in S1 shedding. As expected, the single mutation S621P based on BA.2.86 improved the fusion activity and the mutation P621S in XBB.1.5 dramatically decreased its fusion efficiency (Figure 5E). In line with functional observations, the mutation P621S facilitates formation of an α-helix in the 630 loop, a key modulator for fusion ^1,44^, that would be adopted as a partially disordered loop in P621 variants, to some extent structurally impeding structural rearrangements for subsequent fusion (Figure 5F). These data indicate that the N354 glycosylation coupled with P621S alters multiple virological characteristics, in which cell-cell fusion activity renders the N354 glycosylated variants difficult to be neutralized by antibodies.

### N354 glycosylation specially escapes a subset of ADCC antibodies

Major selective pressures for previous VOCs, such as Delta, BA.2, BA.5 and XBB causing waves of infections globally, came from specific-classes of antibodies driven immune evasion ^3^. Compared to FLip and other XBB variants, BA.2.86 did not show substantial humoral immune escape, while JN.1 with one additional mutation (L455S) on BA.2.86 became more immune-evasive due to extensive resistance across three types of antibodies ^45^. Previously, we determined the escape mutation profiles and epitope distribution of a total of 3051 antibodies isolated from vaccinated or break through infection (BTI) individuals by deep mutational scanning (DMS), which were classified into 12 subgroups (Figure 6A). Immune evasion pattern assays revealed that BA.2.86 sublineages specifically escaped A2, D3, part D4 and many E antibodies when compared to XBB.1.5 (Figure 6B). Strikingly, acquisition of the N354 glycosylation by the K356T substitution largely inactivated group E1, E2.1 and E2.2 antibodies, although these antibodies displayed relatively low but broad neutralizing activities (Figures 6B and S6A). Class E antibodies from E1 to E3 target epitopes on the RBD ranging from left flank through chest to right flank, and most E1, E2.1 and E2.2 antibodies extensively associate with K356 and N354, which has been validated by complex structures, including S309 (E1) (Figures 6C and S6B) ^46^. The mutation K356T could decrease charge/hydrophilic interactions and the N354 glycan fatally induced steric clashes, disabling the binding of most E1, E2.1 and E2.2 antibodies (Figure S6B). Fc-dependent effector mechanisms, e.g., antibody-dependent cell cytotoxicity (ADCC) mediated by natural killer cells, could facilitate viral clearance from infectious individuals. Remarkably, we observed efficient E antibodies-mediated ADCC of SARS-CoV-2 S-transfected cells (Figures 6D and S6C), revealing that the N354 glycosylated variants specially escapes one subset of ADCC antibodies. Together with improved cell-cell fusion, these possibly make the N354 glycosylated variants difficult to be cleared from individuals infected with virus.

**Figure 6:**
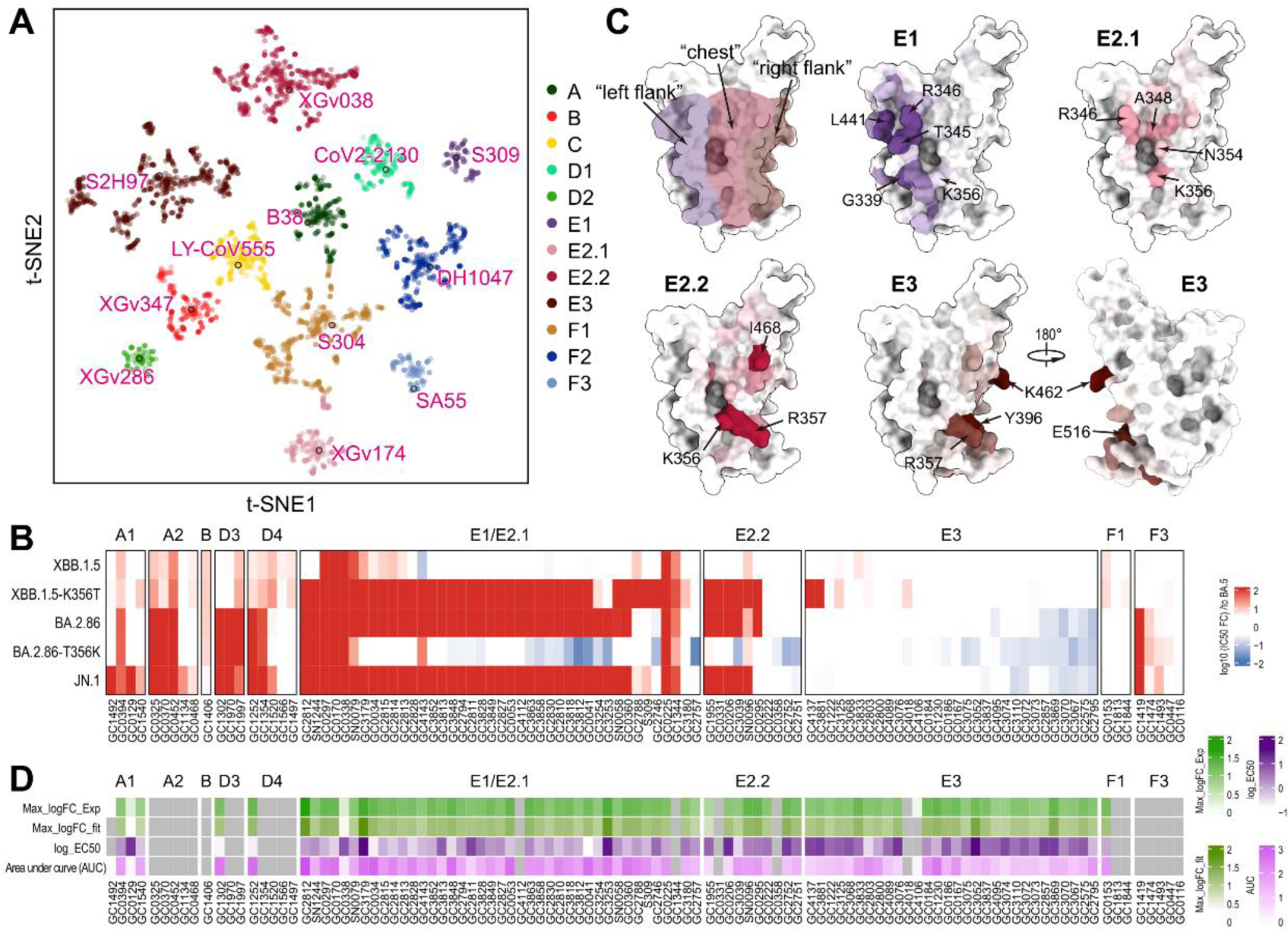
N354 glycosylation specially escapes a subset of ADCC antibodies. (A) t-SNE and unsupervised clustering of antibodies that bind SARS-CoV-2 RBD. Twelve epitope groups were identified from DMS dataset (3051 antibodies). (B) Heatmap of neutralizing activity against XBB.1.5, XBB.1.5-K356T, BA.2.86, BA.2.86-T356K and JN.1 of representative antibodies from 10 epitope groups, relative to BA.5. (C) Mapping of escape scores for antibodies from epitope group E1 (“left flank”), E2.1 (“chest”), E2.2 (“chest”) and E3 (“right flank”) on SARS-CoV-2 RBD (PDB: 6M0J). (D) Heatmap of ADCC effect of RBD antibodies. Four types of color bars represent the base 10 logarithm of the maximum of experiment curve, the base 10 logarithm of the maximum of the fitting curve by 4 parameters fitting, the base 10 logarithm of EC50 and area under curve from Figure S6C.

### N354 glycosylation reduces immunogenicity in hybrid immunity background

In addition to immune escape, viruses generally evolve to acquire new glycosylation sites on the protein surface, a natural phenomenon of glycan shielding, which alters their glycoproteins immunogenicity ^47^. To investigate if the N354 glycosylation may affect its immunogenicity, we first assessed humoral immune responses in naive (non-immunized) BALB/c mice following two-dose primary series immunization with variant S proteins (Figure 7A). All S proteins contained six proline substitutions (S6P) and mutations in the PBCS to stabilize them in the prefusion conformation ^13^. Groups of mice (n = 10 per group) were inoculated intramuscularly with 10 μg of variant S proteins, including BA.5, XBB.1.5, EG.5.1, BA.2.86 and BA.2.86-T356K, on days 0 and 14, and sera were collected at day 28 (2 weeks after the second dose). Administration of BA.5, XBB.1.5 and EG.5.1 S proteins exhibited very low serum 50% neutralizing titers (NT_50_) against BA.2.86, BA.2.86-T356K and JN.1 (using vesicular stomatitis virus-based pseudovirus), meanwhile immunization of BA.2.86 and BA.2.86-T356K resulted in quite limited neutralizing titers against Omicron sublineages, suggesting a large antigenic distance between Omicron and BA.2.86 from single immunity background analysis (Figures 7B and 7C). Notably, the N354 glycosylation decreased BA.2.86 immunogenicity by approximately 40% in comparison with BA.2.86-T356K, rendering BA.2.86 a relatively lower immunogenicity among SARS-CoV-2 variants (Figure 7C).

**Figure 7:**
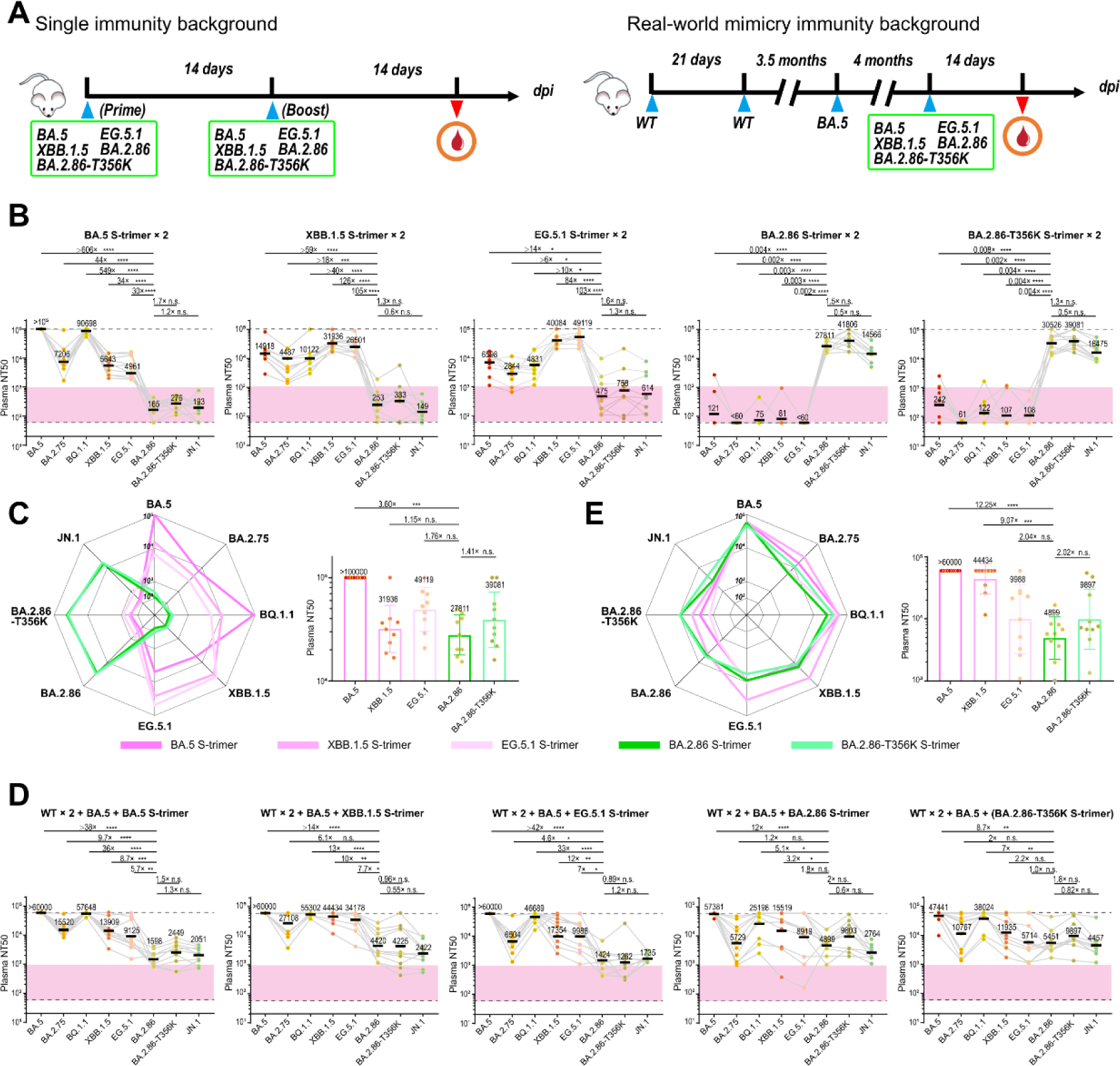
N354 glycosylation reduces immunogenicity in hybrid immunity background. (A) Two cohorts of mice evaluating the immunogenic of various SARS-CoV-2 variants. One cohort consisted of non-immunized BALB/c mice that received two doses of spike proteins (BA.5, XBB.1.5, EG.5.1, BA.2.86, BA.2.86-T356K), with a 14-day interval between each dose. The other cohort mimicked a real-world immunity background, where BALB/c mice were immunized with an inactivated vaccine (two doses of WT + one dose of BA.5) in addition to a single dose of spike protein (BA.5, XBB.1.5, EG.5.1, BA.2.86, BA.2.86-T356K). Blood samples were collected 14 days after immunization. (B) The 50% neutralizing titer (NT50s) against Omicron variants (BA.5, XBB.1.5, EG.5.1, BA.2.86, BA.2.86-T356K) in plasma from non-immunized BALB/c mice background (B) and from BALB/c mice simulating a real-world immune background (D). The p-values were calculated via a two-tailed Wilcoxon signed-rank test for paired samples. Radar plots of the spectrum of neutralization and bar charts of the immunogenicity of the 5 types of immunogens from single immunity background (C) and real-world mimicry immunity background (E).

To further evaluate the effects of the N354 glycosylation in SARS-CoV-2 immune imprinting induced by breakthrough infections, we modeled a real-world mimicry immunity background in mice. To accomplish this, two doses of 0.3 μg CoronaVac (1/10 human dose, an inactivated vaccine derived from WT) were used as primary immunization, then one dose of 0.3 μg inactivated BA.5 vaccine was administrated at 3.5 months after the second dose to mimic BA.5 BTI, and one dose of 10 μg variant S protein at 4 months after the third dose was used to mimic BTI + reinfection (Figure 7A). Compared to single immunity background, single-dose administration of Omicron BA.5, XBB.1.5 and EG.5.1 S proteins under hybrid immunity background displayed ∼5-20-fold improved cross neutralization against BA.2.86 sublineages and single-dose immunization of BA.2.86 and BA.2.86-T356K could produce ∼50-200-fold increased neutralizing titers against Omicron subvariants (Figures 7D and 7E), suggesting existence of hybrid immune imprinting facilitates cross-reactive B cell recall and shortens antigenic distance. As ongoing evolution, an intrinsic trend in gradually decreased immunogenicity for variant S proteins was observed and acquisition of the N354 glycan further induced a ∼2-fold reduction in the immunogenicity under hybrid immunity background, consequently conferring alleviated immune imprinting (Figure 7E). Nonetheless, one-dose booster of BA.2.86, in particular BA.2.86-T356K, under real-world mimicry of immunity background could elicit high levels of neutralizing antibodies against BA.2.86 sublineages, including the currently prevalent JN.1 (Figure 7E), revealing that immune responses can be fine-turned to the BA.2.86 sublineages by boosting with a tweaked (BA.2.86-based) vaccine. These indicate an altered evolution trajectory towards more sophisticated adaptation in humans through acquisition of the N354 glycan.

## Discussion

A selectively favorable mutation spreading all or part of the way through the population generally causes a decrease in the level of sequence variability at nearby genomic sites ^48^, which can be manifested as a selective sweep signature. By using OmegaPlus ^49^ and RAiSD ^50^, we mapped putative sweep regions in 184,224 SARS-CoV-2 genomes deposited in the past 4 months (from 1^st^ Sep 2023 to 1^st^ Jan 2024) from GISAID EpiCoV database. Four similar selective sweep regions were detected in the S from both datasets regardless of whether wild type or BA.2 or BA.5 or XBB was used as a reference (Figure 1D). Two non-synonymous changes (A1067C and A1114G) within the codons for residues 356 (K**→**T) and 372 (T**→**A) of RBD were centrally located in one of the sweep regions, leading to acquisition and loss of N354 and N370 glycosylation, respectively (Figure 1D). Loss of the N370 glycosylation has been shown to be an important evolutionary event for SARS-CoV-2 emergence from animal reservoirs and the enhanced human-to-human transmission during the early stages of the pandemic ^10,11^. Our findings suggest that the N354 glycosylation acquired by variants during the course of the prolonged SARS-CoV-2 pandemic likely confers selective advantage for optimal adaptation in humans through shift in tropism with adjustable infectivity, reduced immunogenicity and elimination-escaped immune evasion.

The conformational dynamics of RBD, and modulation thereof, would render sarbecoviruses cunning to balance host cell attachment and immune escape. The transition to the up state exposures RBD for the binding to hACE2 and is also a prerequisite for S-mediated viral fusion, directly correlating with the infectivity. Thus S proteins from most circulating SARS-CoV-2 variants have been observed in the RBD-up state with a reasonable proportion (>50%). Remarkably, however, recently prevalent BA.2.86 sublineages dominate their S protein in the RBD-down state up to 100% for JN.1 due to acquisition of the N354 glycosylation, shielding RBD from neutralizing antibodies and preventing RBD-hACE2 engagement. These observations suggest a viral evolution trade-off by compromising infectivity in exchanging for a greater evasion of immunity and lower immune imprinting for a goal of co-existence with humans. Surprisingly, the decreased infectivity could be recovered by altered binding mode of HS co-factor to promote the RBD-up conformational transition, apparently through an allosteric mechanism, conferring an adjustable infectivity and a shift in tropism towards HS-abundant cells.

During the process of viral evolution, viruses develop different glycosylation modifications, yielding appreciable impacts on the survival, transmissibility and fitness. In general, the majority of N-glycan adding mutants show decreased infectivity and transmission efficiency ^51^, in turn, immune-shielding glycans are beneficial for immune evasion, which reflects a sophisticated and balanced evolution strategy for N-glycan site accumulation. Further evidence for this has been documented in the viral evolution of Influenza A with additional N-glycan sites every 5–7 years ^52^. Whether the limited glycan shield density observed on SARS-CoV-1, SARS-CoV-2 and Middle East syndrome coronaviruses (MERS) is correlated to the zoonosis of the pathogens is unknown. Notably, among *betacoronavirus* genus, seasonal human coronaviruses HKU1 and OC43 have long co-existed with humans and possess 26-31 N-glycan sites per S monomer, versus 22-23 N-linked glycan sequons in SARS- and MERS-CoVs (Figure S1B). Remarkably, N-glycan sites on OC43 S were accumulated in the past 60 years with approximately 2 N-glycan sites added every 20 years (Figure S1C). A marginal trend in the relationship between N-glycan sites and prevalent time in humans was also observed in HKU1 presumably due to its first isolation and identification in 2004. It’s tempting to speculate that adequate prevalent time might be required to monitor the glycan shield accumulation or HKU1 has evolved to enter a relatively mature stage, bearing ∼30 and 5 N-glycan sites in S monomer and RBD, respectively (Figure S1C). Even so, N-glycan modifications of coronavirus S proteins do not constitute a *bona fide* and effective shield, when compared to the glycan dense of other viruses such as HIV, influenza and Lassa, which may be reflected by overall structure, sparsity, oligomannose abundance and immune evasion ^53^. Although it’s difficult to directly compare viruses in terms of immunogenic responses, SARS-CoVs readily elicit robust neutralizing antibodies that target S proteins following infection or immunization ^54,55^. In contrast, the effective glycan shield of HIV hinders to produce sufficient immune responses and broadly neutralizing antibodies ^56^. We speculated that the high plasticity of SARS-CoV-2 spike RBD may limit the accumulation of glycans on itself. The biological importance of the N354 glycosylation in modulation of SARS-CoV-2 infectivity and immune responses could be applied in coronavirus vaccine research.

## Acknowledgments

We thank Dr. Xiaojun Huang, Dr. Boling Zhu, Dr. Lihong Chen and Dr. Xujing Li for Cryo-EM data collection at the Center for Biological Imaging (CBI) in the Institution of Biophysics, CAS. We also thank Dr. Yuanyuan Chen, Bingxue Zhou and Zhenwei Yang for technical support on surface plasmon resonance (SPR). Work was supported by Ministry of Science and Technology of China (CPL-1233 and SRPG22-003), National Key Research and Development Program (2018YFA0900801), CAS (YSBR-010) and the National Science Foundation Grants (12034006, 32325004 and T2394482). Xiangxi Wang was supported by National Science Fund for Distinguished Young Scholar (No. 32325004) and the NSFS Innovative Research Group (No. 81921005).

## Authors Contributions

X.W., Y.C., R.K.G., Y.W. and B.M. designed the study; P.L., C.Y., B.M., T.X., S.Y., S.L., F.J., Q.Z., Y.Y., Y.R., P.W., Y.L., J.W., X.M. and F.S. performed experiments; P.L., C.Y. and Q.Z. prepared the cryo-EM samples and determined the structures; all authors analyzed data; X.W., Y.C., and R.K.G. wrote the manuscript with input from all co-authors.

---

**Figure S1.**
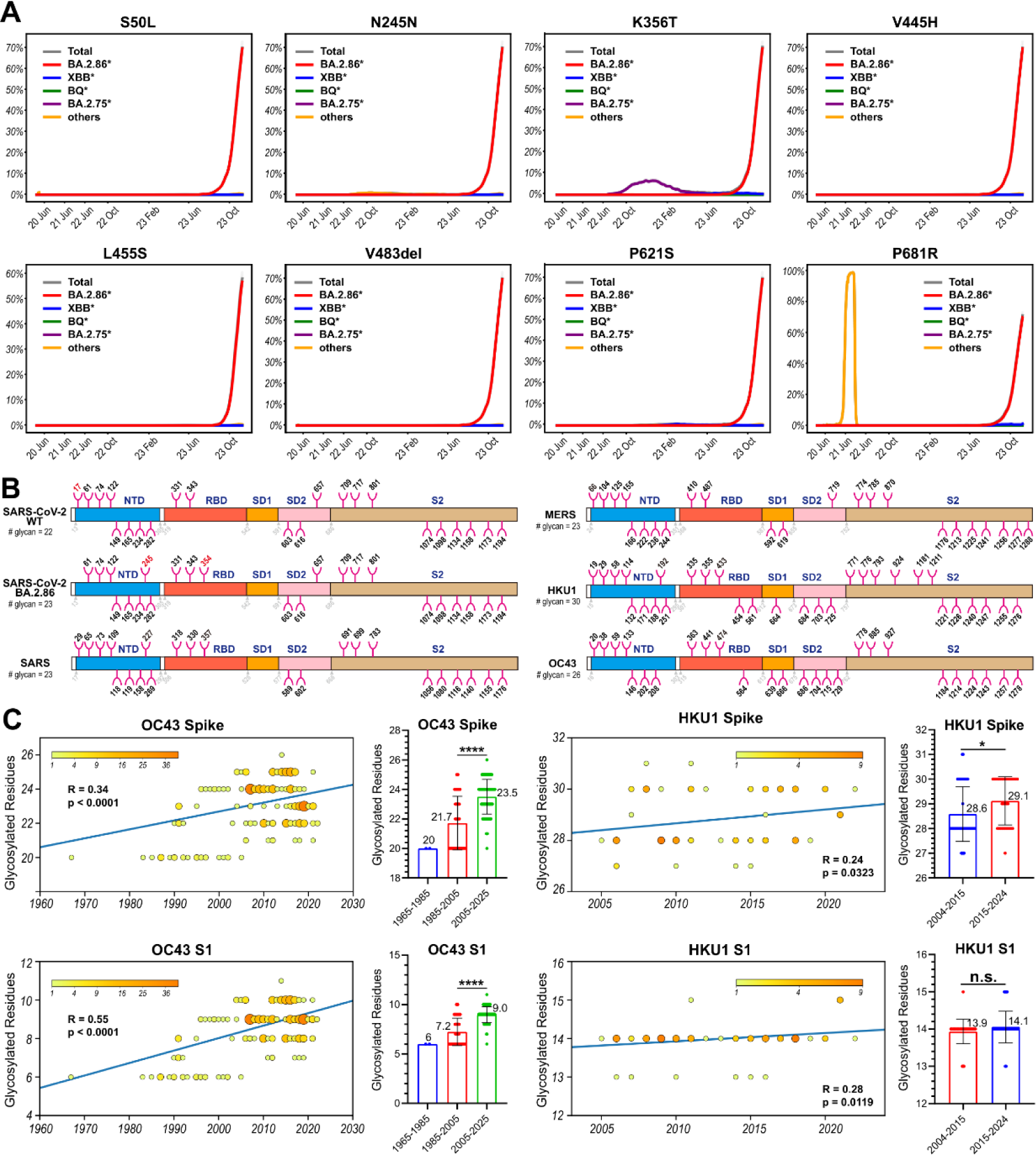
Proportion of typical mutations carried by BA.2.86 spike and distribution of coronavirus glycosylation residues on spike, related to Figure 1. (A) Proportion of SARS-CoV-2 spike residue mutations S50L, H245N, K356T, L455S, V455H, V483del, P621S and P681R from January 2020 to January 2024. (B) A schematic diagram of sequence glycosylation site on SARS-CoV-2 WT, SARS-CoV-2 BA.2.86, SARS, MERS, HKU1 and OC43 spike predicted by “NXS/T” rule. (C) Correlation between the number of glycosylated residues on Spike and S1 over time for OC43 and HKU1.

**Figure S2.**
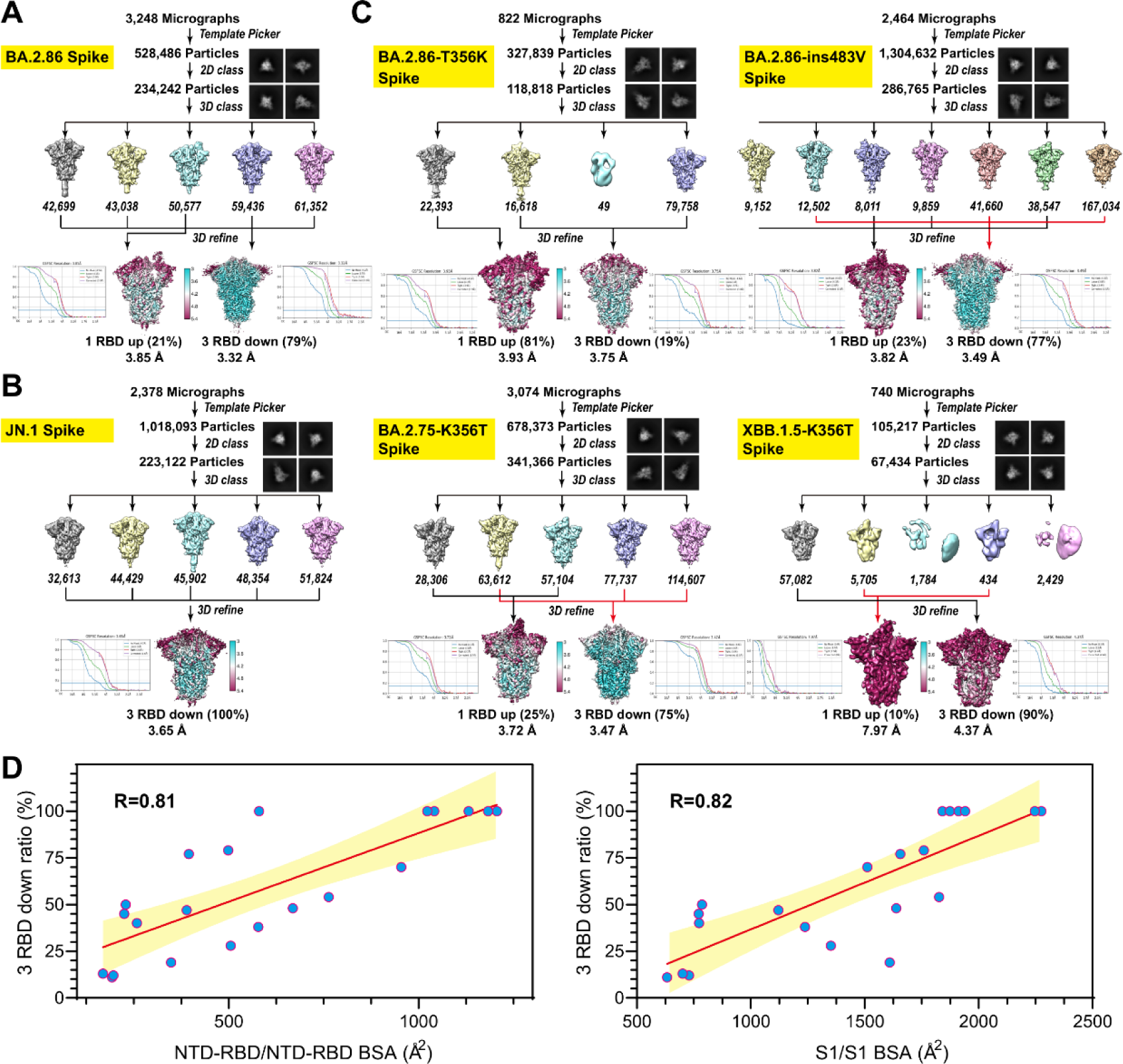
Cryo-EM structures of S-trimer of 6 SARS-CoV-2 variants and linear regression analysis of “RBD down” conformation ratio to buried surface areas (BSA), related to Figure 2. Flow charts, FSC curves and local resolutions of Cryo-EM structure BA.2.86 S-trimer (A), JN.1 S-trimer (B), BA.2.86-T356K S-trimer, BA.2.86-ins483V S-trimer, BA.2.75-K356T S-trimer and XBB.1.5-K356T S-trimer (C). (D) Correlation plots of ratio of spike with “3 RBD down” conformation with NTD-RBD/NTD-RBD BSA and S1/S1 BSA.

**Figure S3.**
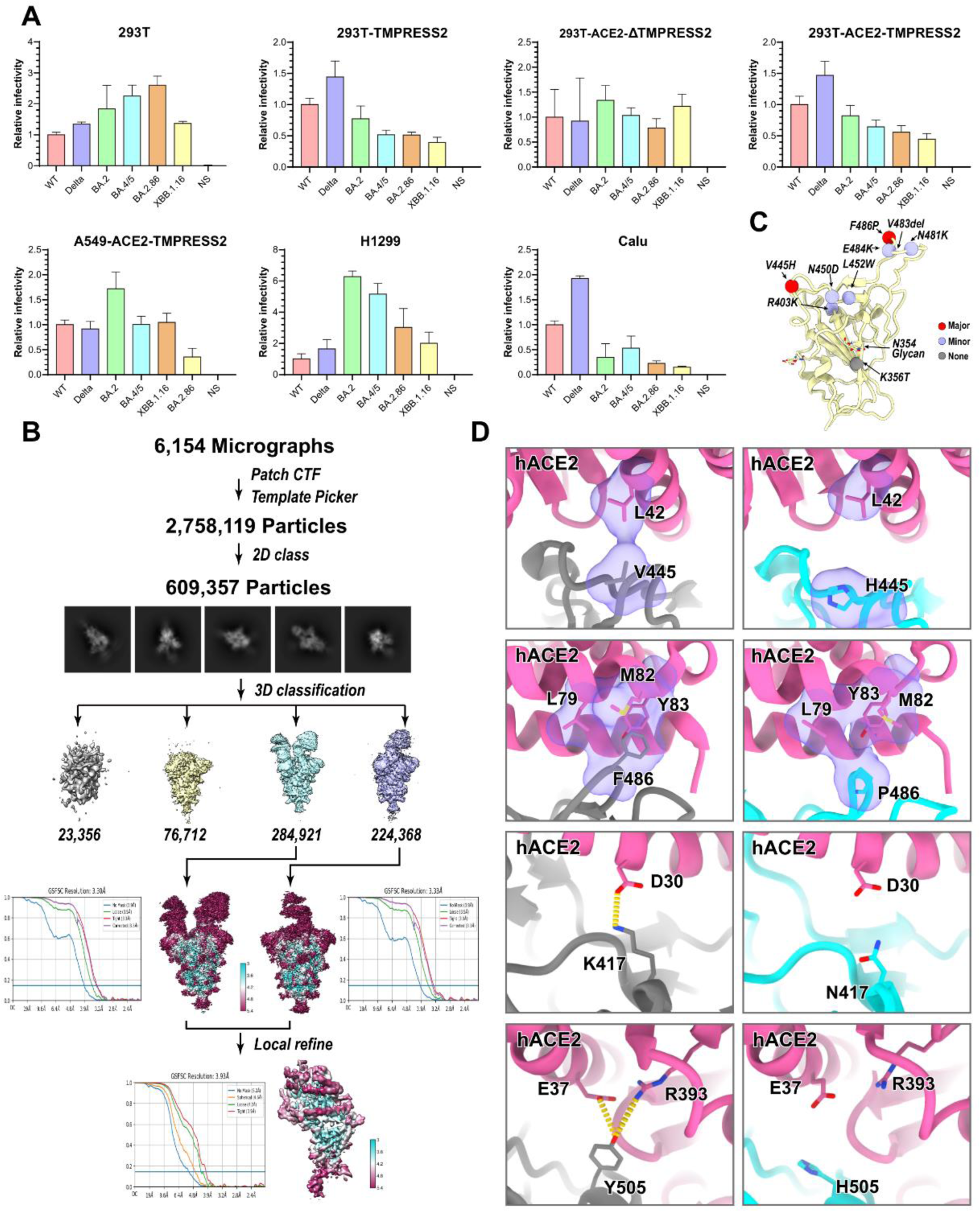
Relative infectivity of SARS-CoV-2 variants in 7 cell lines, cryo-EM structure of BA.2.86 S-trimer in complex with hACE2 and interface details between hACE2 and RBD, related to Figure 3. (A) Normalized SARS-CoV-2 variants pseudovirues entry in HEK293T cells (293T), HEK293T TMPRSS2-overexpressing cells (293T-TMPRSS2), HEK293T cells overexpressing ACE2 and deleted for TMPRSS2 (293T-ACE2-Δ TMPRSS2), HEK293T cells overexpressing ACE2 and TMPRSS2 (293T-ACE2-TMPRSS2), A549 cells overexpressing ACE2 and TMPRSS2 (A549-ACE2-TMPRSS2), H1299 lung cells and Calu-3 lung cells. Error bars represent the mean ± SD of three replicates. (B) Flow chart, FSC curve and local resolution of Cryo-EM structure of BA.2.86 S-trimer in complex with hACE2. (C) Locations of residues on BA.2.86 RBD that play a major (red), minor (light purple) and no (gray) role in binding affinity to hACE2. (D) Structure interpretation of mutation H445V, P486F, N417K and H505Y on BA.2.86 RBD increasing the binding affinity to hACE2. Residues associated with affinity change are shown as sticks. Hydrophobic network is highlighted in light purple and hydrogen bonds are presented as yellow dashed lines. Oxygen atoms are colored in red and nitrogen atoms are colored in blue.

**Figure S4.**
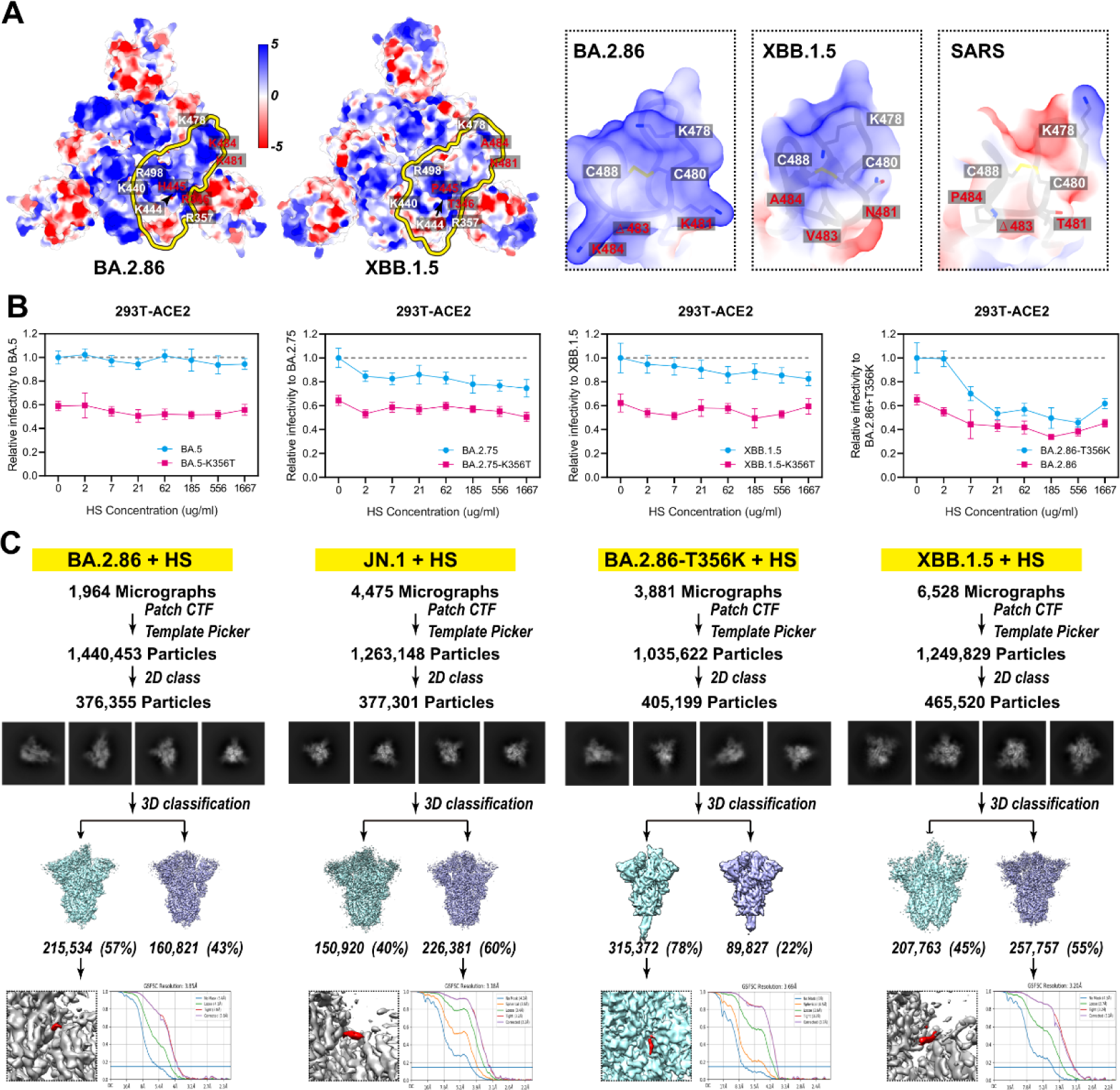
Electrostatic surface of S1 subunit, infectivity with HS treatment and Cryo-EM structure of S-trimer bound to HS, related to Figure 4. (A) Electrostatic surface of BA.2.86 (left) and XBB.1.5 (right) S1 subunit. Yellow circle marked a single RBD. Key residues are labeled and diverse residues are highlighted in red. Electrostatic surface of motif 2 on RBD of BA.2.86, XBB.1.5, and SARS are zoomed in. (B) BA.5, BA.2.75, XBB.1.5, BA.2.86-T356K and their corresponding K356T mutant VSV-based pseudoviruses entry HEK293T cells overexpressing ACE2 (293T-ACE2) treated with various concentrations of free HS. Error bars represent the mean ± SD of three replicates. (C) Flow charts and FSC curves of Cryo-EM structure of S-trimer of BA.2.86, JN.1, BA.2.86-T356K and XBB.1.5 in complex with HS. The density belonging to HS are highlighted in red.

**Figure S5.**
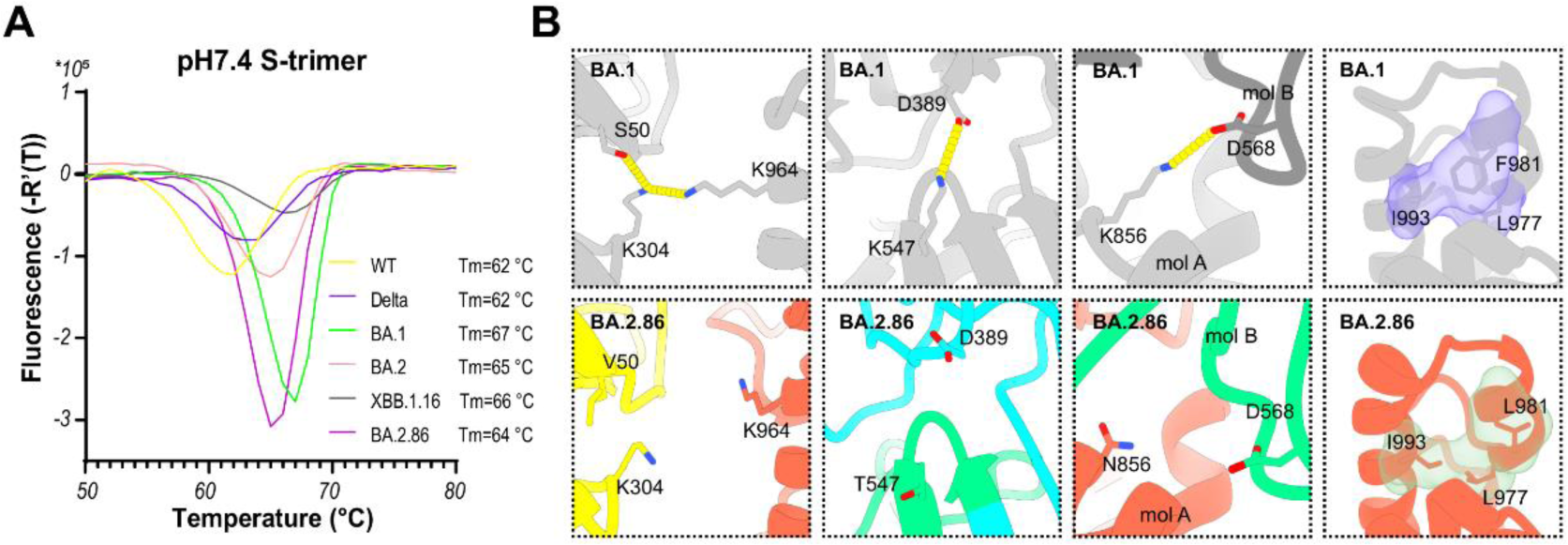
Thermal stability analysis of S-trimer of SARS-CoV-2 variants, related to Figure 5. (A) Thermal stability of WT, Delta, BA.1, BA.2, XBB.1.16 and BA.2.86 S-trimer measured by ThermoFluor Assay at neutral pH. (B) Zoomed-in view of the inter- and intra-subunits of S-trimer interaction details of BA.1 (top) and BA.2.86 (bottom). The residues involved in the interactions are shown as sticks. The hydrogen bonds are shown as yellow dashed lines and hydrophobic network is highlighted in light purple and light green. Different monomers on the same S-trimer are defined as mol A and mol B, respectively.

**Figure S6.**
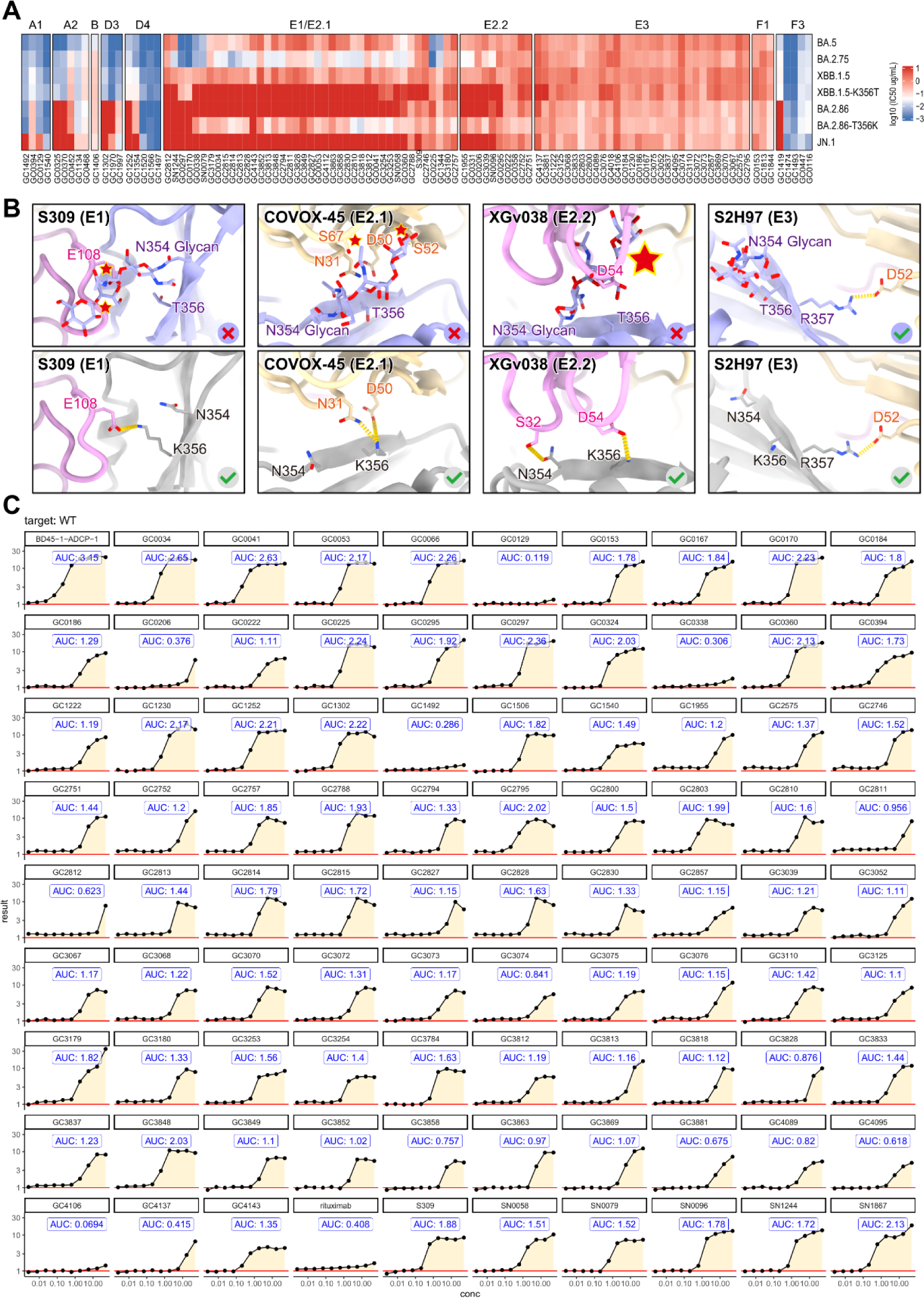
Pseudovirus neutralization assay, structural interpretation of the evasion of antibodies and ADCC assay of antibodies, related to Figure 6. (A) Heatmap of log10 IC50 of antibodies from A1, A2, B, D3, D4, E1, E2.1, E2.2, E3, F1 and F3 epitope groups against BA.5, BA.2.75, XBB.1.5, XBB.1.5-K356T, BA.2.86, BA.2.86-T356K and JN.1 pseudovirus. (B) Cartoon representation of RBD with N354 glycosylation (top) and without N354 glycosylation (bottom) in complex with antibodies S309 (E1), COVOX-45 (E2.1), XGv038 (E2.2) and S2H97 (E3). The key residues of RBDs and antibodies participating interactions are shown as sticks. Atom clashes are shown as red star. The hydrogen bonds are shown as yellow dashed lines. For color scheme, RBD with and without N354 glycosylation are colored in light gray and gray, respectively. Light and heavy chain of antibodies are colored in light yellow and light pink, respectively. (C) The raw data of antibodies ADCC.

**Figure S7.**
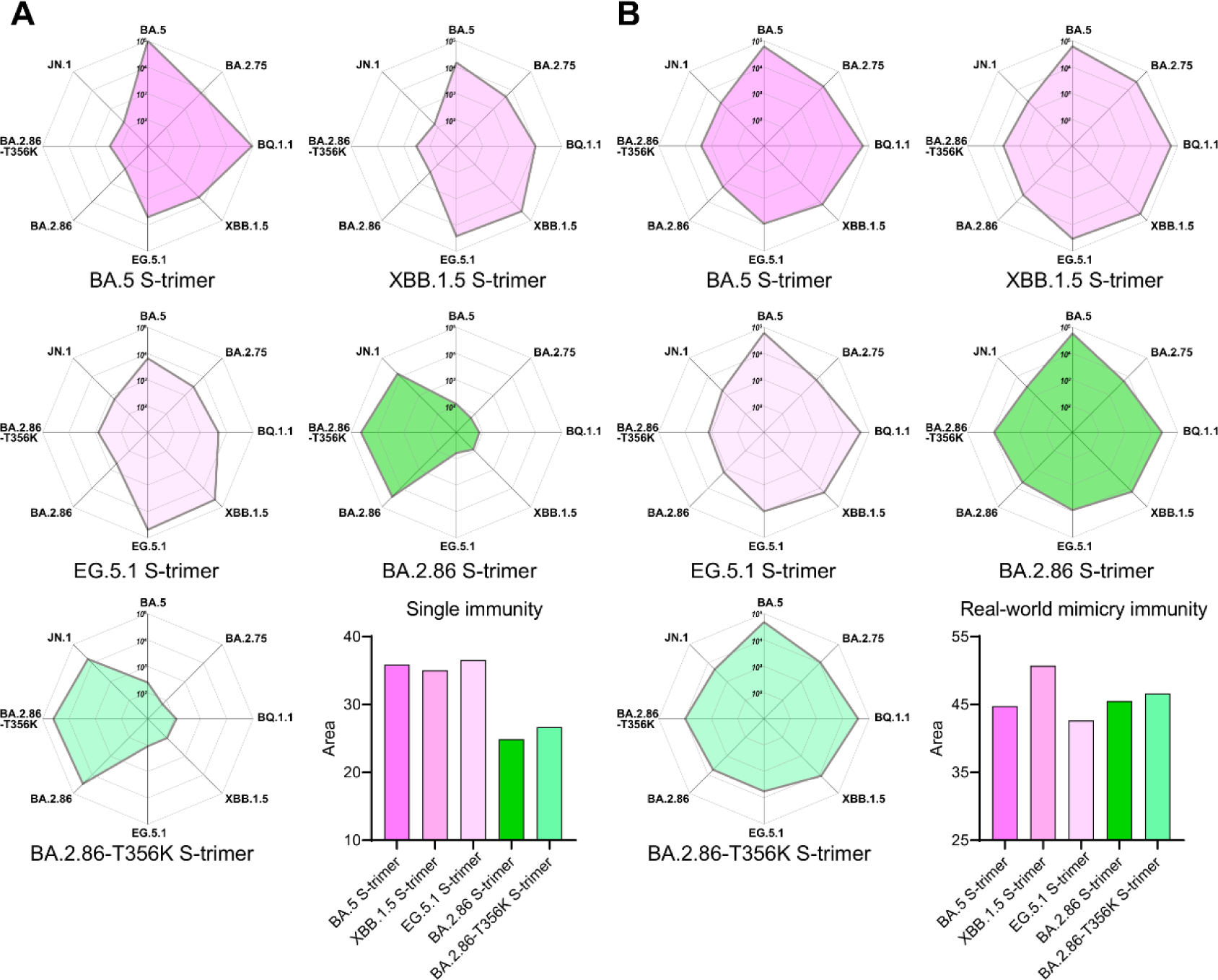
Immunogenicity of BA.2.86 relative to BA.5, XBB.1.5, EG.5.1, and BA.2.86-T356K S-trimer, related to Figure 7. Radar plot of immunogenicity of BA.5, XBB.1.5, EG.5.1, BA.2.86 and BA.2.86-T356K S-trimer under single immunity background (A) and real-world mimicry immunity background (B). The serum neutralizing titers (NT_50_) against 8 pseudovirus (BA.5, BA.2.75, BQ.1.1, XBB.1.5, EG.5.1, BA.2.86 and BA.2.86-T356K) are log10-scaled to generate the radar plot and to calculate the area of the radar plot, which is displayed by bar chart. Color schemes are consistent with figure 7.

**Table S1.**
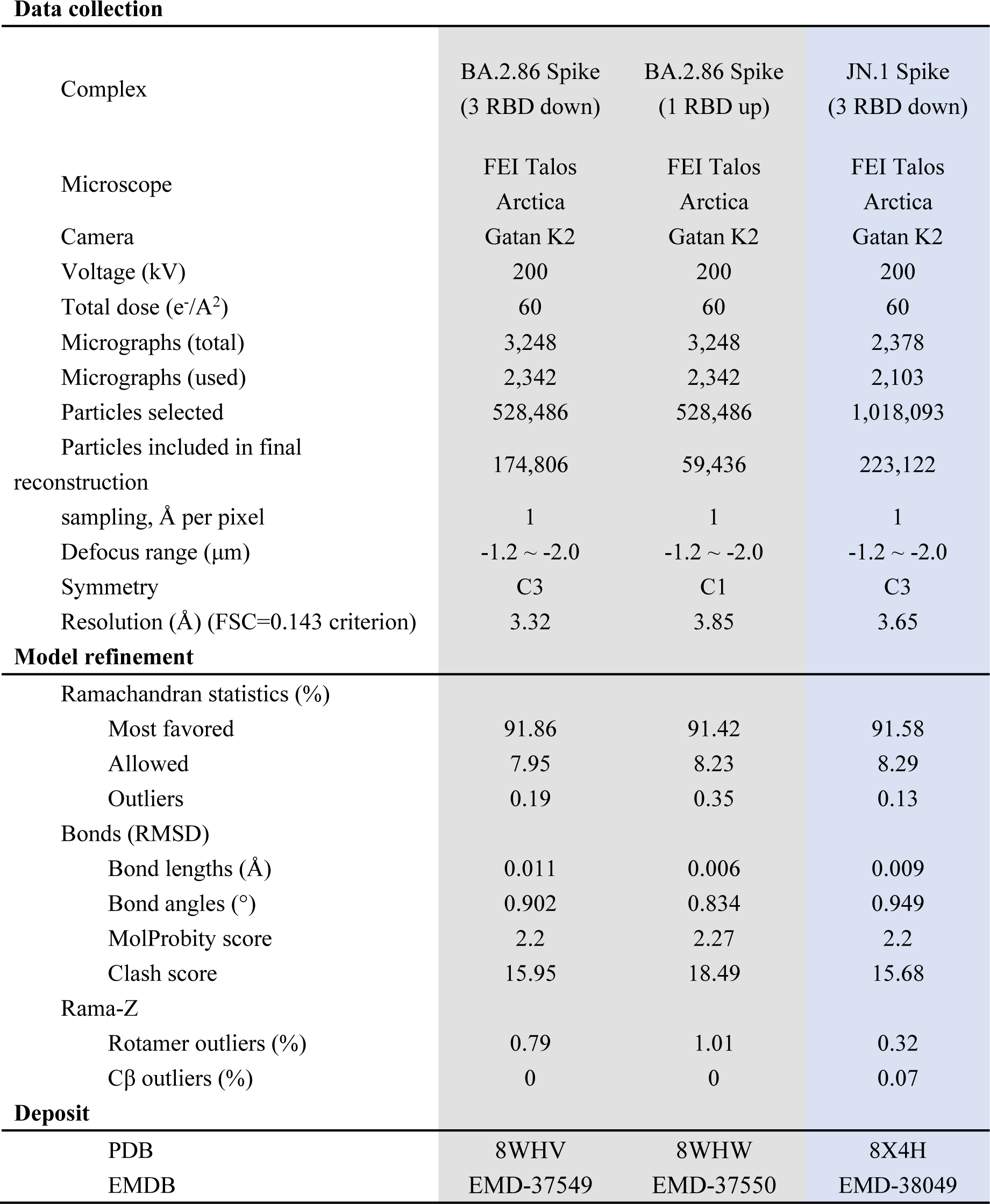
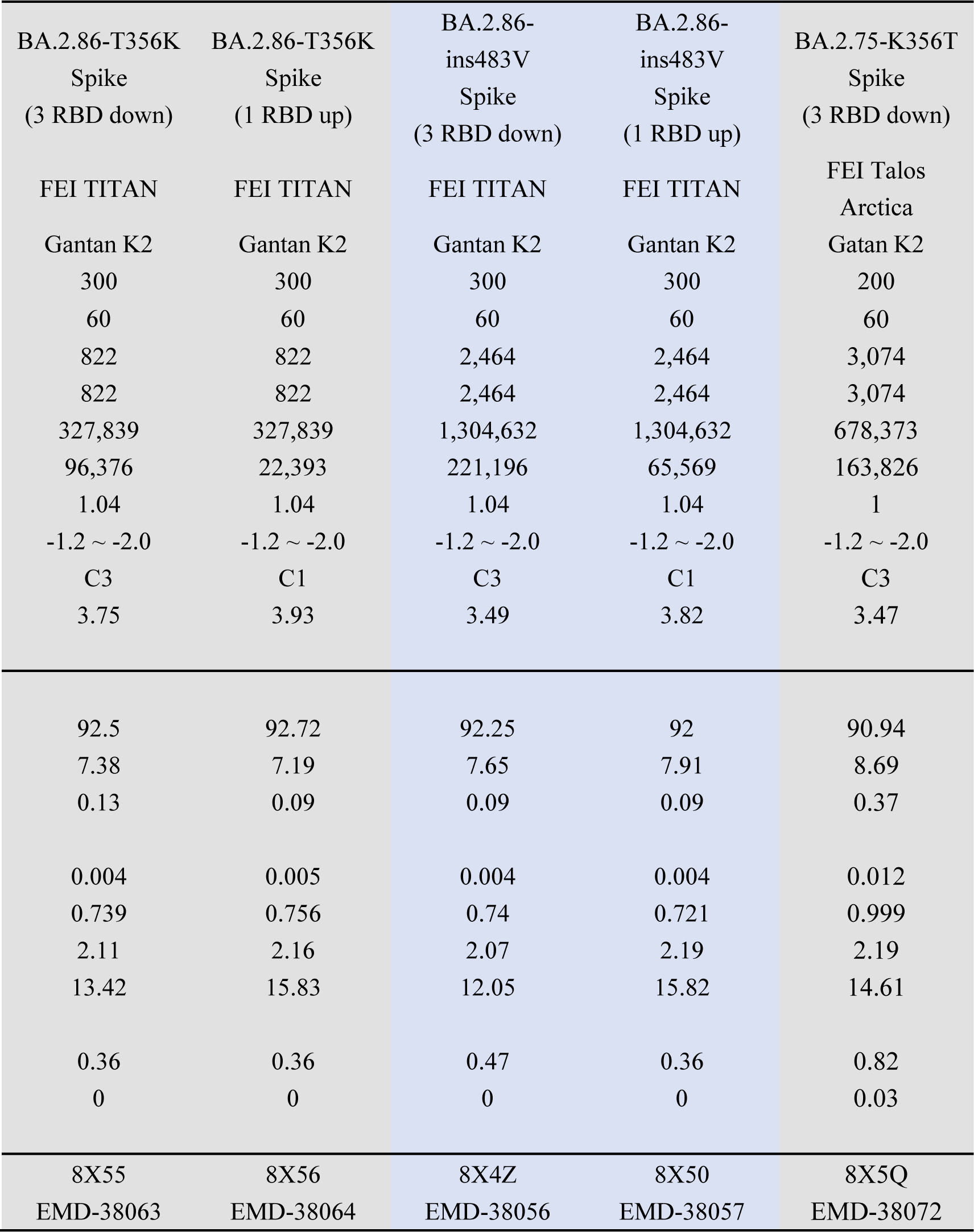
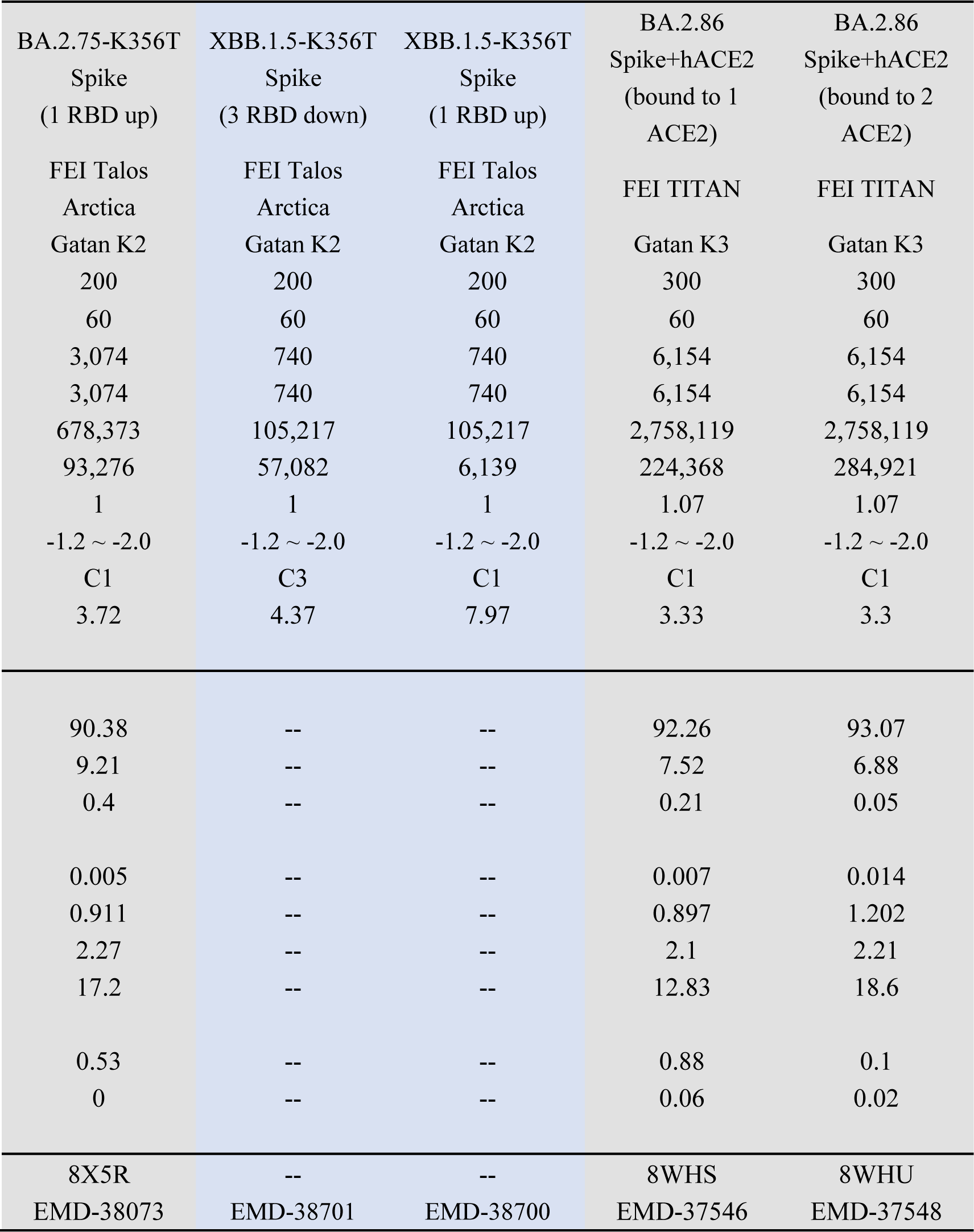
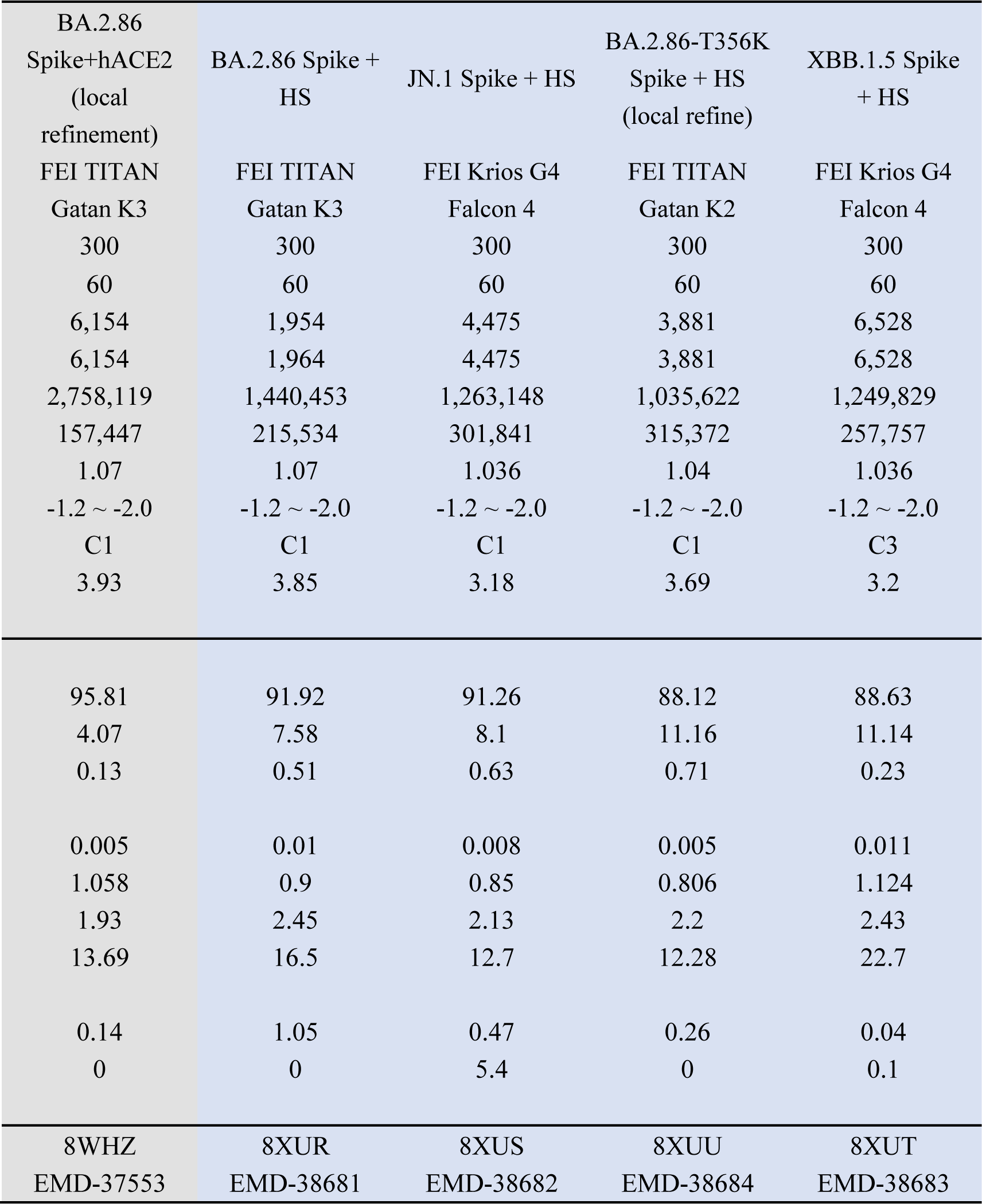
Cryo-EM data collection, processing, and validation statisticss.

## STAR METHOD

### KEY RESOURCE TABLE

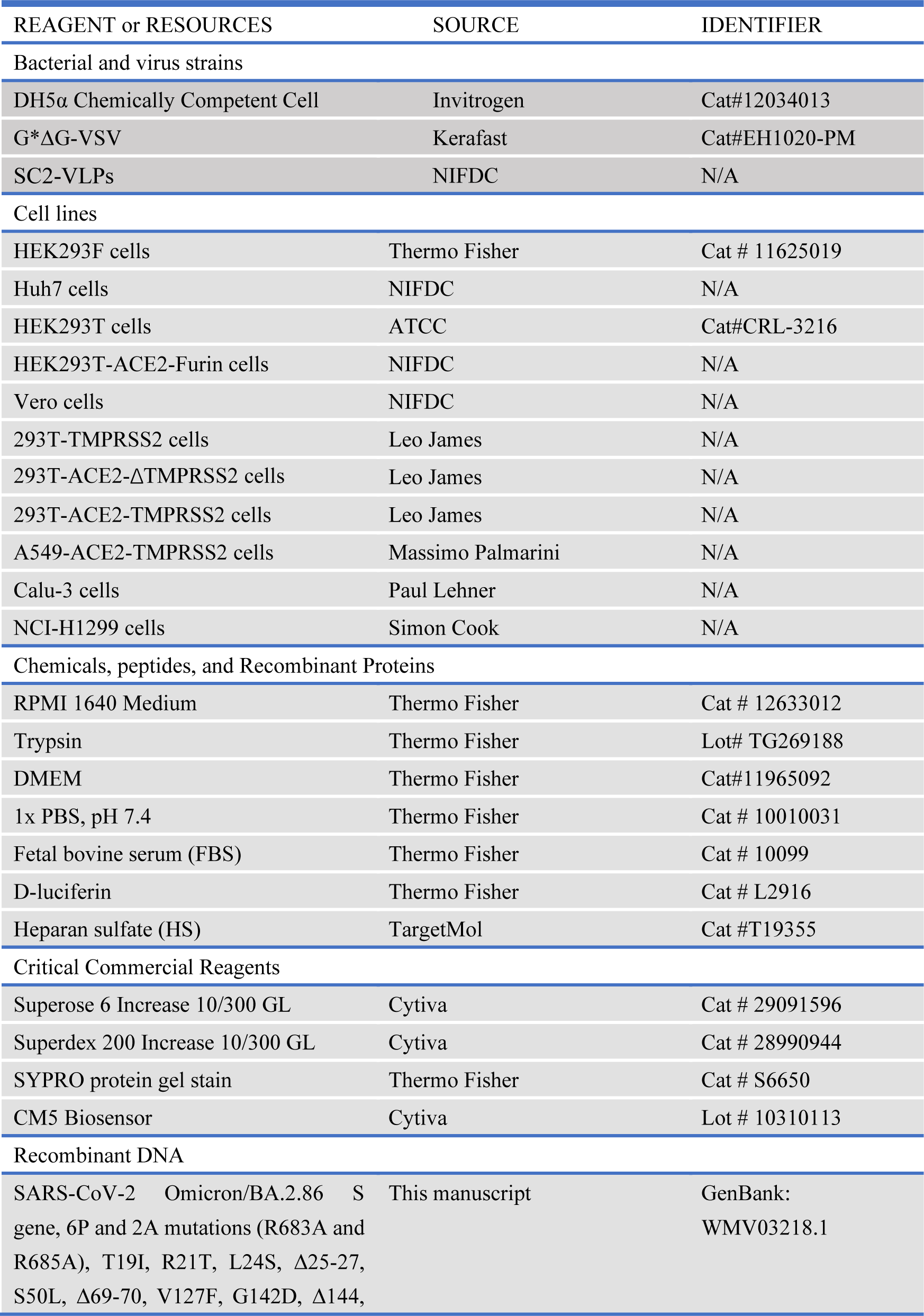
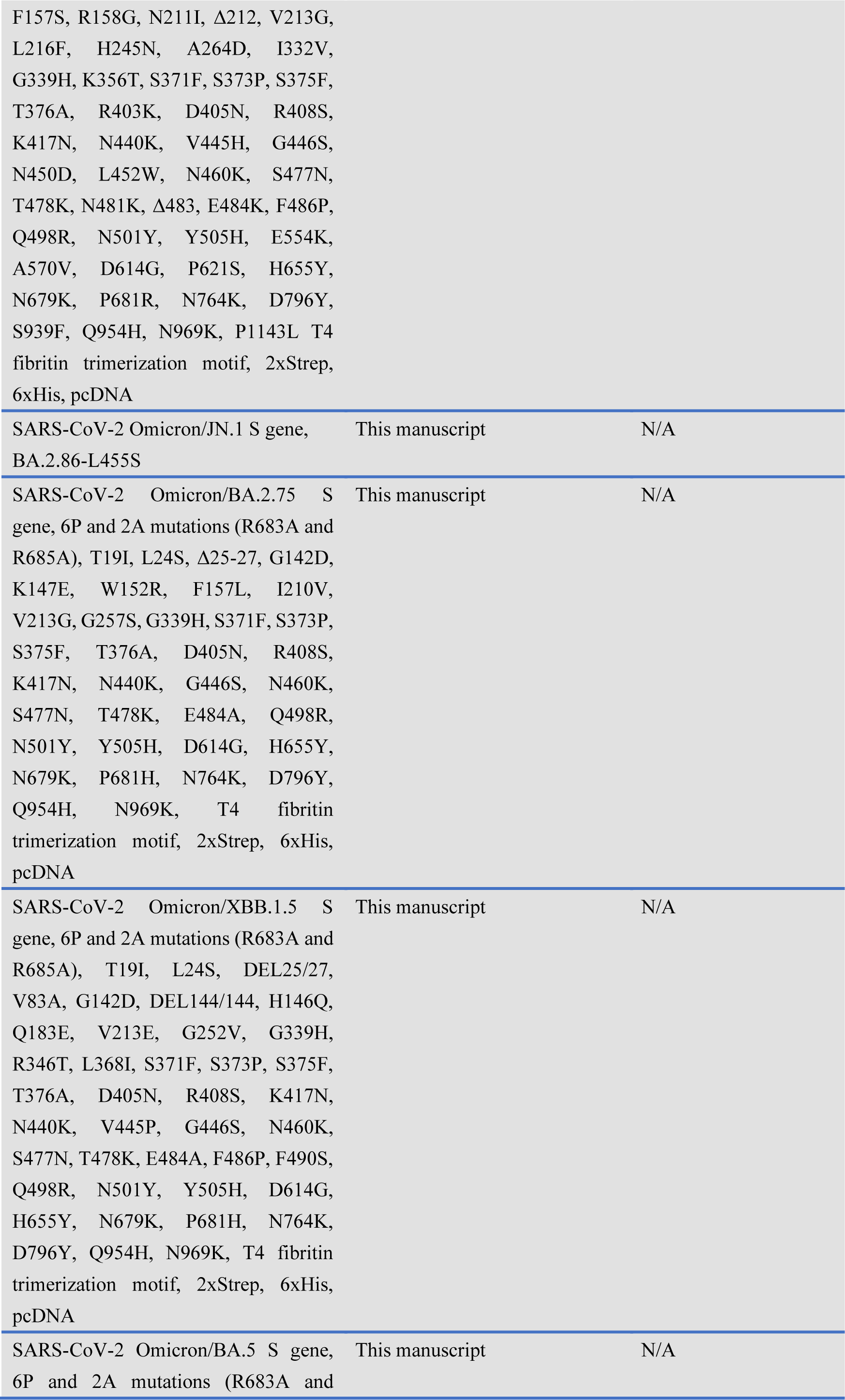
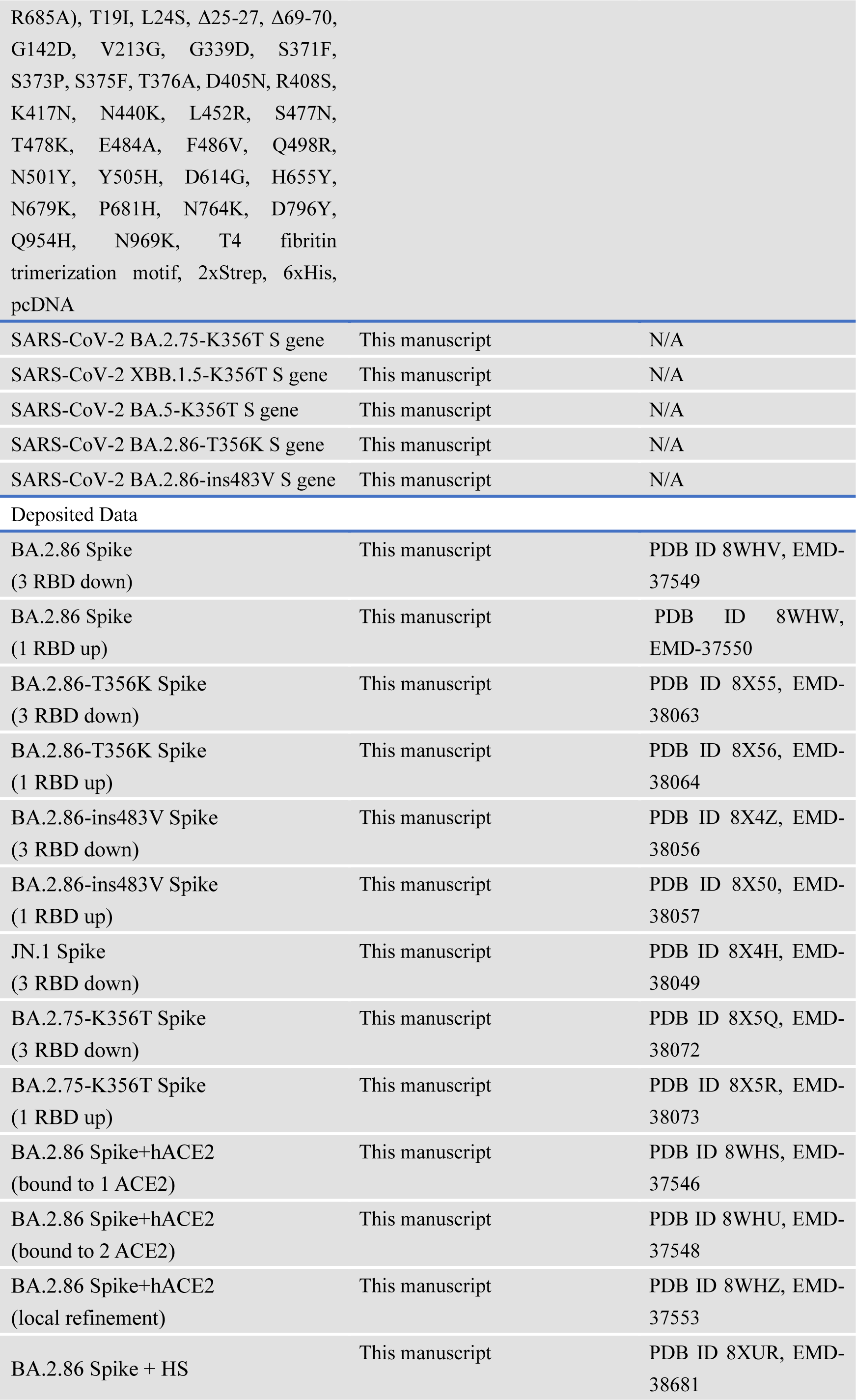
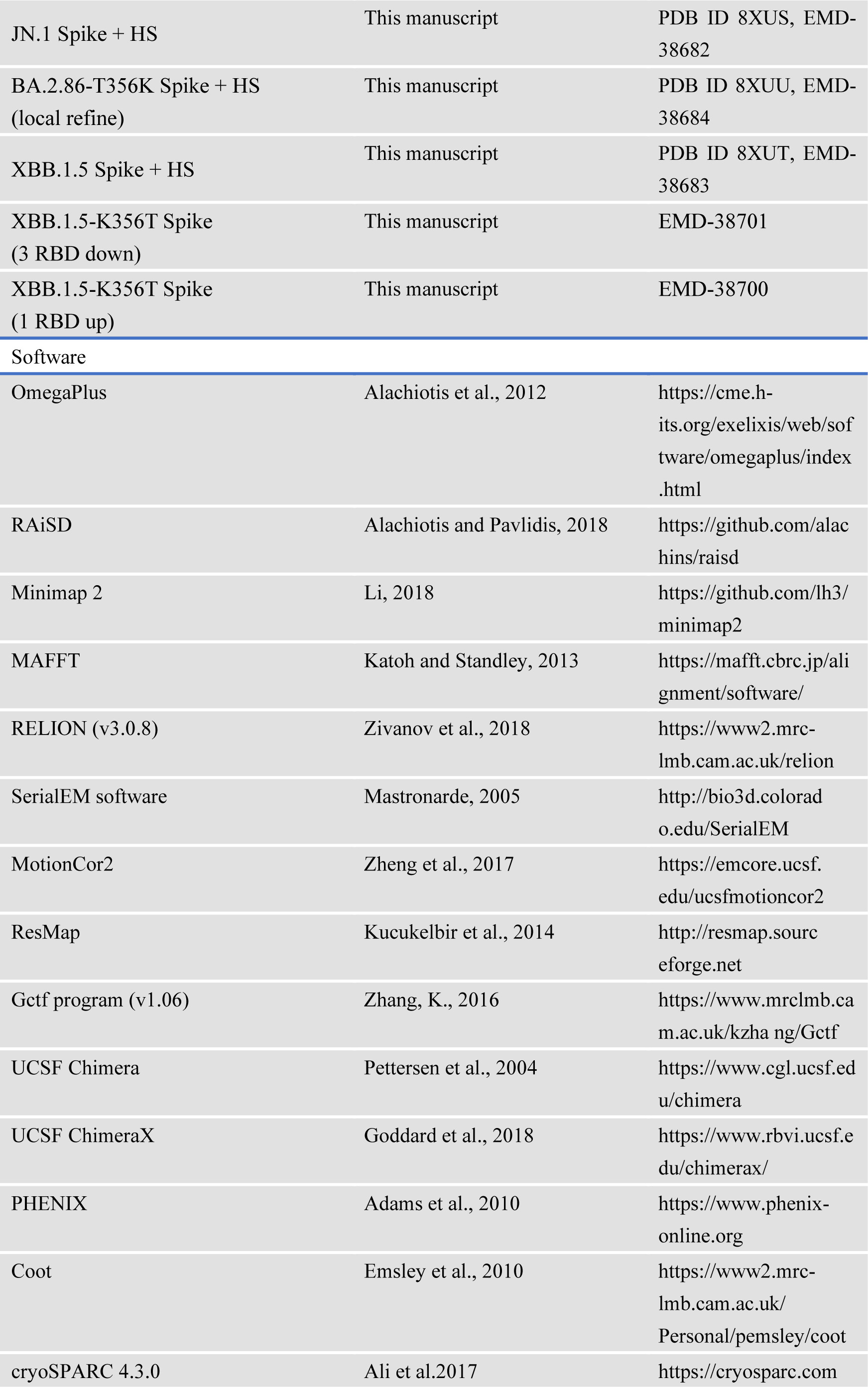
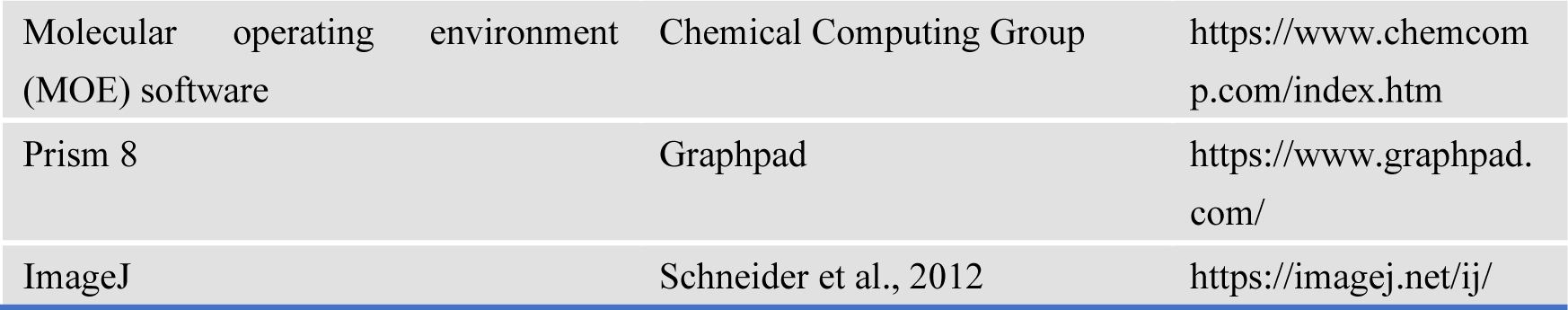

#### RESOURCE AVAILABILITY

##### Lead contact

Further information and requests for resources and reagents should be directed to and will be fulfilled by the Lead Contact, Xiangxi Wang

##### Materials Availability

All plasmids generated in this study are available from the Lead Contact with a completed Materials Transfer Agreement.

##### Data and code availability

The density maps of the BA.2.86 S-trimer with 3 RBDs down conformation, BA.2.86 S-trimer with 1 RBD up conformation, JN.1 S-trimer with 3 RBDs down conformation, BA.2.86-T356K S-trimer with 3 RBDs down conformation, BA.2.86-T356K S-trimer with 1 RBD up conformation, BA.2.86-ins483V S-trimer with 3 RBDs down conformation, BA.2.86-ins483V S-trimer with 1 RBD up conformation, BA.2.75-K356T S-trimer with 3 RBDs down conformation, BA.2.75-K356T S-trimer with 1 RBD up conformation, BA.2.86 S-trimer bound to 1 hACE2, BA.2.86 S-trimer bound to 1 hACE2, BA.2.86 S-trimer bound to 2 hACE2s, local refinement of BA.2.86 S-trimer in complex with hACE2, BA.2.86 S-trimer bound to HS, JN.1 S-trimer bound to HS, BA.2.86-T356K S-trimer bound to HS and XBB.1.5 S-trimer bound to HS have been deposited in the Electron Microscopy Data Bank under the accession codes of EMD-37549, EMD-37550, EMD-38049, EMD-38063, EMD-38064, EMD-38056, EMD-38057, EMD-38072, EMD-38073, EMD-37546, EMD-37548, EMD-37553, EMD-38681, EMD-38682, EMD-38684 and EMD-38683, respectively. The corresponding structural coordinates have been deposited in the Protein Data Bank: 8WHV, 8WHW, 8X4H, 8X55, 8X56, 8X4Z, 8X50, 8X5Q, 8X5R, 8WHS, 8WHU, 8WHZ, 8XUR, 8XUS, 8XUU and 8XUT. For XBB.1.5-K356T S-trimer with 3 RBDs down conformation and XBB.1.5-K356T S-trimer with 1 RBD up conformation, only density maps have been deposited in the Electron Microscopy Data Bank under the accession codes of EMD-38701 and EMD-38700.

### EXPERIMENTAL MODEL AND SUBJECT DETAILS

#### Cell lines

HEK293F cells were cultured in StarFect 293 Transient Transfection Medium. HEK293T and its derivative cell lines including 293T-ACE2ΔTMPRSS2, 293T-ACE2-TMPRSS2, 293T-TMPRSS2 were cultured in Dulbecco’s Modified Eagle’s Medium (DMEM) supplemented with 10% fetal bovine serum (FBS). The cultures were maintained at 37 ℃ in an incubator supplied with 5% CO_2_. Calu-3 cells were maintained in Eagle’s minimum essential medium containing 10% FBS and 1% PS. Vero cells, Huh-7 cells, A549-ACE2-TMPRSS2 were maintained in Dulbecco’s modified Eagle’s medium (DMEM) containing 10% FBS and 1% PS. NCI-H1299 cells were maintained in RPMI containing 10% FBS and 1% PS.

#### SARS-CoV-2 pseudovirus

The SARS-CoV-2 pseudovirus was constructed as previously described using VSV pseudotyped virus (G*ΔG-VSV) and VLP pseudotyped virus (SC2-VLPs). For VSV pseudotyped virus, D614G, BA.2, BA.2-K356T, BA.2-P621S, BA.5, BA.5-K356T, BA.2.75, BA.2.75-K356T, XBB.1.5, XBB.1.5-K356T, XBB.1.5-P621S, EG.5.1, BA.2.86, BA.2.86-T356K, BA.2.86-S621P and JN.1 were constructed and used. For VLP pseudotyped virus, BA.5, BA.5-K356T, BA.2.75, BA.2.75-K356T, XBB.1.5, XBB.1.5-K356T, BA.2.86 and BA.2.86-T356K were constructed and used.

### METHOD DETAILS

#### Putative Selective Sweep Region Detection

Genomic scans for selective sweeps were performed by two programs. The one is OmegaPlus v3.0.3 ^49^ and the other is RAiSD v2.9 ^50^. A total of 184,224 SARS-CoV-2 spike protein sequences were retrieved from the GISAID EpiCov database (https://www.gisaid.org/) from Sep.01,2023 to Dec.31,2023. Sequence reads were aligned to SARS-CoV-2 WT (NCBI Reference Sequence/NC_045512.2), BA2 (BA.2_hCoV-19/France/IDF-IPP08725/2022|EPI_ISL_10071318), BA5 (BA.5_hCoV-19/South_Africa/NCV1255/2022|EPI_ISL_12587877), XBB (XBB_hCoV-19/USA/NY-NYULH8854/2022|EPI_ISL_15427610) reference using Minimap2 ^57^. Sequences with aligned lengths less than 3200 were excluded from the analysis, leaving 163,500 sequences for alignment using MAFFT ^58^. OmegaPlus was performed with the following parameters: SARS-CoV-2 spike sequence was divided into 1000 bins (-grid 1000). The calculation of linkage disequilibrium values between SNPs utilized a defined window range, specifically set at 20bp as the minimum and 200bp as the maximum (-minwin 20-maxwin 200). RAiSD was performed with the following parameters: The grid size to specify the total number of evaluation points was set to be 1000 (-G 1000); Missing data imputation was enabled (-M 1; per SNP). The sliding window size was set to be 200bp (-w200). Combination statistics of both programs, the top 20% intersecting regions detected were selected as the candidate sweep reigons (-COT 0.2).

#### Protein expression and purification

The full-length sequence information of Spike (S) from BA.2.86 was obtained from the NCBI (GenBank: WMV03218.1). The BA.2.86 spike and RBD protein genes were obtained by using the BA.2.75 gene as a template and performing overlapping PCR. The JN.1, BA.2.86-T356K, BA.2.86-ins483V Spike and JN.1, BA.2.86-T356K, BA.2.86-N354Q, BA.2.86-K403R, BA.2.86-D450N, BA.2.86-H445V, BA.2.86-W452L, BA.2.86-K481N, BA.2.86-ins483V, BA.2.86-K484A, BA.2.86-P486F, BA.2.86-N417K, BA.2.86-H505Y, BA.2.86-(N417K+H505Y), BA.2.86-(L455F+F456L), BA.2.86-(N417K+H505Y+L455F+F456L) RBD protein genes were obtained by using the BA.2.86 Spike and RBD genes as a template and performing overlapping PCR. The BA.2.75-K356T Spike and RBD protein genes were obtained by using the BA.2.75 gene as a template and performing overlapping PCR. The XBB.1.5-K356T Spike and RBD protein genes were obtained by using the XBB.1.5 gene as a template and performing overlapping PCR. To improve protein expression and stabilize the trimeric conformation, proline substitution was performed at residues 817, 892, 899, 942, 986, and 987 in all Spike gene constructs. And all spikes were modified to incorporate 2A mutations (R683A and R685A). Additionally, the C-terminus of the constructs was modified by adding the T4 fibritin folding domain. To facilitate protein purification, His or Strep II tags were attached at the C-terminus of all gene constructs. The spike and RBD proteins were obtained using a eukaryotic expression system. Plasmids containing the target protein were transiently transfected into suspended HEK293F cells and cultured at 37°C in a constant-temperature shaker with 8% CO_2_ for 72 hours. After collecting the cell supernatant, preliminary purification was performed using Ni-NTA or affinity StrepTactin resin chromatography. The proteins were further purified using Superdex 200 10/300GL (Cytiva) or Superose 6 10/300 (Cytiva) in phosphate-buffered saline (PBS) at pH 7.4 to obtain high-purity proteins.

#### Surface Plasmon Resonance

Surface plasmon resonance (SPR) was utilized for quantifying the binding affinity between the antigen and receptor, as well as the antigen and heparan sulfate (HS). In investigating interactions between the receptor and antigen, human ACE2 (hACE2) was immobilized as the stationary phase, while the SARS-CoV-2 RBDs acted as the mobile phase. For evaluating the affinity between HS and RBDs, the RBDs were immobilized as the stationary phase, and HS molecules were used as the mobile phase. These experiments were conducted at a temperature of 25°C, employing the Biacore8K biosensor on the S series CM5 chip (Cytiva) for data detection and recording. The raw data curves were analyzed and fitted using the Biacore 8K evaluation software (GE Healthcare) employing a 1:1 binding model.

#### Determination of Spike stability

The stability of spike protein trimers of WT, Delta, BA.1, BA.2, BA.2.86, and XBB.1.16 at neutral pH (pH = 7.4) was evaluated using the ThermoFluor Assay. 5 μg sample of the spike protein was added to a 25 μL reaction system containing a final concentration of 1× SYPRO Orange dye (Invitrogen, USA) as a fluorescence probe. The protein was heated from 25°C to 99°C at a rate of 1°C/min using the QuantStudio™ 6 Flex Real-Time PCR System instrument (Applied Biosystems, USA), and changes in fluorescence signals were recorded during this temperature gradient. Data analysis and curve plotting were performed using GraphPad Prism 9.4.0 (GraphPad Software Inc.).

#### Cryo-EM sample preparation and data collection and model building

Purified Spike trimer protein samples from SARS-CoV-2 variants BA.2.86, JN.1, BA.2.86-T356K, BA.2.86-ins483V, BA.2.75-K356T, and XBB.1.5-K356T were diluted to a concentration of 1.0 mg/mL in PBS buffer, pH 7.4. Similarly, to prepare the spike/hACE2 complex sample, the BA.2.86 Spike protein was mixed with hACE2 at a molar ratio of 1:1.2, and the Spike/HS complex were mixed at a molar ratio of 1:1000, while maintaining a constant concentration of 1.0 mg/mL for the Spike. 3μL of the sample was pipetted onto pre-treated porous carbon-coated gold grid (C-flat, 300 mesh, 1.2/1.3, Protochips Inc.). The Vitrobot (FEI) was operated in a no-force mode to blot the sample for 6 seconds under 100% relative humidity and room temperature conditions. Subsequently, the sample was rapidly plunge frozen into liquid ethane.

Cryo-EM datasets were collected using a 200 kV FEI Krios ARCTICA or 300 kV FEI Titan microscope (Thermo Fisher) equipped with K2, K3, or Falcon 4 detectors. Movies were recorded with 32 frames at an exposure time of 0.2 seconds per frame, resulting in a total dose of 60 e ^−^ Å ^-2^. The automated single-particle data collection using SerialEM resulted in a final pixel size of 1 Å, 1.036 Å, 1.04 Å or 1.07 Å.

Data processing was performed using cryoSPARC (v4.3.0) and Relion (v3.0.8). The data underwent several steps including Motion Correction, CTF Estimation, Create Templates, Template Picker, Extract from Micrographs, 2D classification, 2D selection for Ab-initio Reconstruction, and subsequent Homogeneous Refinement. To enhance the density around the RBD/RBD-ACE2 region, local refinement was conducted using UCSF Chimera (v1.13.1) and CryoSPARC (v3.2.1). Structural modeling and refinement were performed using WinCoot (v0.9.8.1) and Phenix (v1.20.1). Figures were generated using UCSF ChimeraX (v1.6.1).

#### Molecular Docking

Electrostatic potential maps of the BA.2.86 S-trimer and BA.2.86-T356K S-trimer were generated in UCSF ChimeraX (v1.6.1). A heparan sulfate fragment was docked to the BA.2.86 and BA.2.86-T356K RBD using the Molecular operating environment (MOE) software (Version 2020.09). The docking was done with default parameters.

#### Infectivity assay

Spike-pseudotyped VSV are prepared as described previously ^59^. The spike genes (D614G, Delta, BA.1, BA.2, BA.5, BA.5-K356T, BA.2.75, BA.2.75-K356T, XBB.1.5, XBB.1.5-K356T, EG.5.1, BA.2.86, BA.2.86-T356K, JN.1) were optimized using mammalian codons and inserted into the pcDNA3.1 vector. Afterwards, the plasmids were transfected into 293T cells using Lipofectamine 3000 (Invitrogen). These cells were separately infected with G*ΔG-VSV pseudotyped virus (Kerafast) and virus-like particles (SC2-VLPs) pseudovirus. After incubation, the supernatant containing the pseudovirus was collected, filtered through a 0.45 μm filter membrane, and stored at −80℃ for future use.

We used HEK293T-hACE2 cells, Vero cells and Huh-7 cells as targets in infectivity assays. After quantification using RT-PCR, 100 ul aliquots of the diluted virus were introduced into individual wells of 96-well cell culture plates. Chemiluminescence monitoring was conducted following a 24-hour incubation period with a temperature of 37°C and a CO_2_ concentration of 5%. The supernatant for each sample was adjusted to a volume of 100 ul to ensure consistency. A mixture of luciferase substrate and cell lysis buffer (PerkinElmer, Fremont, CA) was prepared and added to each well at a volume of 100 ul. 150 ul of the resulting lysate was transferred to opaque 96-well plates after 2 min. 2 PerkinElmer Ensight luminometer was used to detect the luminescence signal, and the data was recorded in terms of relative luminescence unit (RLU) values. Each experimental group consisted of two replicates and the entire set of experiments was repeated three times. For infectivity assay related to heparin sulfate (HS), Virus-like particles (SC2-VLPs) were selected to infect HEK293T cells overexpressing ACE2 and Furin (293T-ACE2/Furin) treated with various concentrations of free heparin sulfate (HS).

#### Western blot analysis

Cell-cell fusion assays were described previously ^60^. At 48 h of infection, cells and culture medium were collected. The culture media were centrifuged and the supernatants were collected. After that, an equal volume of clarified supernatants was mixed with 20% PEG6000 in PBS and centrifuged at 12,000g for 30 min at 4 °C, followed by pellet resuspension in 1× SDS sample buffer.

For cell lysates, the collected cells were washed and lysed in lysis buffer (Cell Signalling) and the lysates were diluted with 4 × sample buffer (Bio-Rad) and boiled for 10 min before analysed using western blotting. The following antibodies were used for protein detection: mouse anti-SARS-CoV-2 S1 antibodies (MAB105403, R&D systems), rabbit anti-SARS-CoV-2 S monoclonal antibodies (PA1-41165, Thermo Fisher Scientific), horseradish peroxidase (HRP)-conjugated anti-rabbit and anti-mouse IgG polyclonal antibodies (Cell Signalling). The ChemiDoc Touch Imaging System (Bio-Rad) was used to detected Chemiluminescence. The cleavage ratio of S1 or S2 to FL in virions was determined by densitometry using ImageJ software (NIH).

#### Cell-cell fusion assay

Cell-cell fusion assays were described previously ^41^. In brief, HEK293T GFP11 and Vero-GFP1-10 cells were seeded at 80% confluence at a 1:1 ratio in 48-well plates the day before. Cells were co-transfected with 0.5 µg of spike expression plasmids. An Incucyte was used to measure cell–cell fusion and fusion was determined as the proportion of green area to total phase area. To measure cell surface spike expression, HEK293 cells were transfected with S expression plasmids and stained with rabbit anti-SARS-CoV-2 S S1/S2 polyclonal antibodies (Thermo Fisher Scientific, PA5-112048, 1:100). Negative control is normal rabbit IgG (SouthernBiotech, 0111-01, 1:100), and Secondary antibodies are APC-conjugated goat anti-rabbit IgG polyclonal antibodies (Jackson ImmunoResearch, 111-136-144, 1:50). The surface expression level of S proteins was analysed using FACS Canto II (BD Biosciences) and FlowJo v.10.7.1 (BD Biosciences).

#### Antibody expression and purification

SARS-CoV-2 RBD-specific mAbs were synthesized as described previously. Briefly, antibody heavy and light chain genes were synthesized by GenScript, inserted into pCMV3-CH, pCMV3-CL or pCMV3-CK vector plasmids by infusion (Vazyme), and co-transfected into Expi293F cells (Thermo Fisher) using polyethylenimine. Transfected cells were cultured at 36.5°C in 5% CO2 and 175 rpm for 6-10 days. Expression fluid was then collected and centrifuged, and the supernatants containing monoclonal antibodies were purified with Protein A magnetic beads (GenScript). Purified antibodies were verified by SDS-PAGE.

#### Pseudovirus neutralization assay

SARS-CoV-2 variants (BA.5, BA.2.75, BQ.1.1, XBB.1.5, XBB.1.5-K356T, EG.5.1, BA.2.86, BA.2.86-T356K, JN.1) spike-pseudotyped virus was constructed based on a vesicular stomatitis virus (VSV) pseudovirus packaging system, as described previously. Spike gene is inserted into pcDNA3.1 vectors. G*ΔG-VSV virus (VSV G pseudotyped virus, Kerafast) and spike plasmids were transfected to HEK293T cells (American Type Culture Collection [ATCC], CRL-3216). After culture, the pseudovirus in the supernatant was harvested, filtered, aliquoted, and frozen at −80°C for further use.

We used Huh-7 cells (Japanese Collection of Research Bioresources [JCRB], 0403) as targets in pseudovirus neutralization assays. Plasma samples or mAbs were serially diluted in culture media and mixed with pseudovirus, and incubated for 1 h in a 37°C incubator with 5% CO_2_. Digested Huh-7 cells were seeded in the antibody-virus mixture. After 1 day incubation, the supernatant was discarded. D-luciferin reagent (PerkinElmer, 6066769) was added into the plates and incubated in the dark for 2 min, and cell lysis was transferred to plates for detection. The luminescence values were measured by a microplate spectrophotometer (PerkinElmer, HH3400). IC50 values for mAbs and NT50 values for plasma were determined by fitting a logistic regression model.

#### ADCC assays

The full-length spike gene sequence of SARS-CoV-2 (GenBank: MN908947) was synthesized, with mutations for prefusion stabilization and modification on the furin cleavage site (Spike-6P2A or Spike-6P/GSAS). The Spike gene was inserted into the pLVX-puro vector. HEK293T cells(10 million) were transfected with 20.7 μg helper plasmid (pSPAX2), 13.8 μg VSV-G expression plasmid (pMD2.G) and 1 μg full-length spike expression vector (pLVX-puro) using the PEI transfection system(Yeasen, 40816ES03) to generate the lentivirus. Cell supernatant containing lentivirus was collected 48 hours after transfection, centrifuge at 500 g at 4 °C for 10 min and filtered through a surfactant-free cellulose acetate 0.45 mm syringe filter. 5 ml lentivirus was used to infect 1 million low passage HEK293T cells. At 72 hours post infection, cells were stained with SA55-FITC for RBD labeling, single cell clone was sorted into a 96-well plate containing selective medium (DMEM+10 % FBS + 10 μg/ml puromycin + 1 % penicillin-streptomycin solution) using BD Aria II cell sorter in FITC channel. Single clone cell was expanded and tested for the Spike expression level via continuous puromycin selection and flow cytometry cell sortings. The clone with the highest Spike expression level (target cells) for amplification and used to evaluate the ADCC effect of antibodies.

The ADCC effector cells (Human CD16a Jurkat reporter cells) are gift from Youchun Wang, which were engineered to express both the NFAT response element driving luciferase expressing systems and human CD16a receptor with 158V mutation for higher affinity to IgG1 and IgG3 isotypes. Similarly, we used flow cytometry to analyze the expression of human CD16 on the effector cells. Cells were incubated with BD Pharmingen PE Mouse Anti-Human CD16 and subjected for FACS analyses using BD Aria II cell sorter.

To detect antibodies’ potency to mediate ADCC, mAbs were pre-diluted in a reaction medium of 1640 medium (HyClone) containing 10% FBS (Gibco) and 1% penicillin-streptomycin solution. Add serial dilutions of antibodies in 10μL to wells. Meanwhile, add 10μL of reaction medium to the unstimulated control wells. Digest target cells and plate the target cells at a density of 1.67 ×10^6^ cells/ml in 384-well culture plates in 10μL of reaction medium, then aliquots of pre-diluted antibodies were incubated with target cells in 384-well culture plates for 10 minutes at 37 ℃ in the 5% CO2 incubator. Subsequently, add the effector cells (Human CD16a Jurkat reporter cells) at a density of 1.67 ×106 cells/ml in 10μL of reaction medium to the culture wells. The mixtures were further incubated at 37℃ in the 5% CO2 incubator for 18 hours. Finally, add 30μL reagent of Stable-Lite Luciferase Assay System (Vazyme, DD1202) to each well and incubate in the dark for 2 minutes. The chemiluminescence signals were collected by PerkinElmer Ensight. The ADCC luciferase (luc) fold induction was calculated by the fold change of the relative light unit (RLU) values of test wells to control blank wells. The area under curve (AUC, log-concentration v.s. log-fold-change) values are calculated and used as a metric to evaluate the capability of each antibody to mediate ADCC. ADCC assays for all mAbs were conducted in two replicates.

#### Protein vaccine preparation and mouse immunization

The spike proteins, including BA.5, EG.5.1, XBB.1.5, BA.2.86 and BA.2.86-T356K were used for mouse immunization. All of these proteins were purified as previously described.

Animal experiments were carried out under study protocols approved by Rodent Experimental Animal Management Committee of Institute of Biophysics, Chinese Academy of Sciences (SYXK2023300) and Animal Welfare Ethics Committee of HFK Biologics (HFK-AP-20210930). Six- to eight-week-old female BALB/c mice were used for experiments. The mice were kept under a 12-hour light and 12-hour dark cycle, with room temperatures maintained between 20 °C and 26 °C. The humidity levels in the housing area ranged from 30% to 70%. Mice were immunized according to schemes in Fig. 7. Briefly, two cohorts of BALB/c mice were established to evaluate the immunogenic of various SARS-CoV-2 variants. For the first cohort, mice had a pre-existing immune background and received 0.3 ug SARS-CoV-2 WT inactivated vaccine (CoronaVac, against SARS-CoV-2 wild-type) as the primary vaccination at six- to eight weeks age, followed by a booster dose after 14 days. When these mice reached 5.5 months of age, they were given an additional boost using the omicron (BA.5) inactivated vaccine (against SARS-CoV-2 BA.5 variant) with a dose of 0.3 ug as well. For the last dose of this cohort, the mice were intramuscularly administered 10 μg, 1 mg/ml of spike protein from 5 different variants (BA.5, XBB.1.5, EG.5.1, BA.2.86 and BA.2.86-T356K) as immunogen. For the other cohort, mice did not undergo any form of immunization. They were intramuscularly administered 2 doses of 10 ug, 1 mg/ml of 5 types of spike proteins as described before. The interval between the two injections is 14 days. The adjuvants for inactivated vaccines and proteins are Al+CpG. Blood samples were collected on the 14th day post-immunization, and serum was obtained through centrifugation.

### QUANTIFICATION AND STATISTICAL ANALYSIS

The statistical analyses were performed using Prism 8 (GraphPad). All experiments were conducted three times or more, as stated in the figure legends. When comparing two groups, unpaired t-tests were used for statistical evaluation; for multiple comparisons, one-way ANOVA was employed without post hoc correction. Non-linear regression analysis was performed using the inhibitor versus responder least-squares fit approach to calculate IC50 values and confidence intervals. The mean plus standard deviation (SD) values are represented by error bars in the figures. The methods section and figure captions provide a list of the specific statistical tests utilized. Statistical significance was determined according to the following framework: ns: p > 0.05, *: p ≤ 0.05, **: p ≤ 0.01, ***: p ≤ 0.001, ****: p ≤ 0.0001.

